# N6-methyladenosine (m^6^A) RNA modifications control the antiviral response to influenza A virus infection by limiting IFITM2/3 expression

**DOI:** 10.64898/2026.10.01.755989

**Authors:** Piotr Kwiatkowski, Liudmila A. Mikheeva, Holly Digby, Clément R. Bouton, Hannah Jones, Stuart J. D. Neil, Michelle L. Holland, Chad M. Swanson

## Abstract

N^6^-methyladenosine (m^6^A) RNA modifications regulate diverse cellular processes, including the response to virus infection. However, how m^6^A controls antiviral gene expression and whether effects on viral replication are due to direct modification of viral RNA or regulation of cellular transcripts remain unclear. Herein, we analysed how m^6^A modifications control the cell-autonomous innate immune response to influenza A virus (IAV) infection. Depletion or inhibition of the m^6^A writer METTL3 or the m^6^A reader YTHDF2 restricts IAV replication. However, using a quantitative method to identify m^6^A sites, we did not detect substantial m^6^A modification of viral RNA. In contrast, many IAV-induced cellular transcripts were m^6^A modified in their coding region, including those that encode pattern recognition receptors and direct antiviral proteins. Inhibition of interferon signalling restored IAV replication after METTL3 inhibition, highlighting that m^6^A controls the antiviral response to infection. METTL3 inhibition increased expression of the paralogous antiviral proteins IFITM2 and IFITM3, which were required for the antiviral activity of the METTL3 inhibitor. We propose that m^6^A modifications regulate the expression of antiviral proteins to limit the magnitude of their induction and balance antiviral activity with maintaining cellular homeostasis.

## INTRODUCTION

N^6^-methyladenosine (m^6^A) is the most abundant internal modification on eukaryotic mRNA (1–3). Most m^6^A marks on mammalian mRNA are installed by a large multicomponent methyltransferase ‘writer’ complex containing the RNA methyltransferase METTL3 and several auxiliary proteins, including METTL14 (4–7). The m^6^A writer complex co-transcriptionally modifies adenosines in nascent mRNA, selectively targeting a five-nucleotide ‘DRACH’ motif, in which the central A is methylated (D = A, G or U; R = G or A; H = A, C or U) (8–10). Most adenosines in a DRACH motif can be methylated unless they are protected by the exon junction complex (EJC) or other proteins deposited through pre-mRNA splicing in an ∼100 nucleotide window surrounding an exon-exon junction, leading to long internal exons and terminal exons containing most m^6^A sites (11–15). Many genes express transcripts that contain m^6^A modified sites, though the density and modification stoichiometry of these sites is variable (16–18). While m^6^A was initially suggested to be reversible through the activity of the ‘eraser’ proteins FTO and ALKBH5, subsequent work has shown that FTO does not act as an m^6^A demethylase and ALKBH5 has only a minor influence on m^6^A abundance, suggesting that m^6^A is largely irreversible in most contexts (19–22).

m^6^A modifications have been implicated in nearly every stage of post-transcriptional gene regulation, though they are best characterised for promoting mRNA decay (22–27). The effects of m^6^A modification on mRNA are mediated through changes in RNA structure or the action of specific RNA-binding proteins (RBPs) known as m^6^A ‘readers’, which are recruited to the modified transcripts (3,22). The best characterised reader proteins contain a YT521-B homology (YTH) domain, including YTHDC1, YTHDF1, YTHDF2, and YTHDF3 (3,8,22). YTHDC1 resides in the nucleus and regulates post-transcriptional processes such as mRNA splicing or nuclear export (28,29). The YTHDF1-3 proteins are cytoplasmic and predominantly mediate mRNA degradation, potentially through recruitment of the CCR4-NOT deadenylation complex to modified transcripts (23,25–27,30). Recently, YTHDF-mediated RNA decay has been shown to be predominantly co-translational due to ribosomal stalling at m^6^A sites, leading to ribosome collisions and subsequent RNA decay (31,32).

The functional importance of m^6^A-mediated control of gene expression extends to diverse cellular processes, including regulation of development, cell differentiation and stress responses (22). More recently, m6A has been proposed to function as part of an “RNA surveillance” system in which it helps to recognise and eliminate transcripts that are potentially deleterious for cells (33). This model postulates that properly spliced mRNAs are not targeted for m^6^A-mediated decay due to the exon-junction complex and other splicing-associated proteins protecting coding regions from m^6^A modifications, while transcripts with processing errors or those derived from transposable elements or viruses are marked by m^6^A for degradation. Conversely, other models propose that m^6^A marks on viral RNA facilitate efficient replication, suggesting that viruses have evolved to hijack the m^6^A machinery to promote one or more steps of their life cycle (34,35). Therefore, a key question is whether viral RNA contains sites that are m^6^A modified with high stoichiometry and whether modification at these sites have a functional role in viral replication. Importantly, the effects of m^6^A during infection may not be limited to viral RNA, since some cellular transcripts encoding viral cofactors or restriction factors contain m^6^A modifications (36–38). Overall, understanding the roles and relative contributions of m^6^A modifications on viral and cellular transcripts is essential to determine how this modification controls viral replication.

One way in which m^6^A modification of cellular transcripts could influence viral replication is through regulation of the cell-autonomous innate immune response. Recognition of pathogen-associated molecular patterns (PAMPs) by pattern recognition receptors (PRRs), including double stranded RNA (dsRNA) recognition by RIG-I and MDA5, initiates antiviral and pro-inflammatory signal transduction pathways (39). These culminate in the activation of the transcription factors, including NF-κB and IRF3/7, thereby triggering the production of type I interferons (39). Secreted interferons subsequently signal through the IFNAR receptor to activate the JAK-STAT pathway and induce hundreds of interferon-stimulated genes (ISGs) that establish an antiviral state and directly restrict viral replication (40). Emerging evidence suggests that m^6^A can regulate this antiviral programme at multiple stages. At the level of viral RNA sensing, m^6^A modification of viral RNA has been proposed to prevent the recognition of dsRNA by PRRs (35,41). Furthermore, m^6^A has also been implicated in regulating the expression of immune signalling molecules downstream of PRRs (42–45). Transcripts encoding direct antiviral proteins have also been proposed to be regulated by m^6^A (37,38). m^6^A-mediated regulation of antiviral protein expression through RNA decay could therefore provide a mechanism for controlling the magnitude and duration of the antiviral response, allowing effective restriction of viral replication while limiting disruption to cellular homeostasis. Importantly, recent technological advances, such as small molecule inhibitors that enable acute inhibition of METTL3 activity (46) and techniques that allow quantification of m^6^A stoichiometry at specific sites (22) now allow this to be explored in higher resolution.

Among the viruses whose replication is modulated by m^6^A is influenza A virus (IAV), a highly contagious respiratory virus responsible for seasonal epidemics and recurring pandemics (47). The IAV genome consists of eight negative-sense single-stranded viral RNA (vRNA) segments that are transcribed in the nucleus into mRNA for protein synthesis and replicated into complementary RNA (cRNA) for progeny vRNA production (48,49). While the m^6^A writer complex is required for IAV replication, the mechanism underlying this effect remains unclear. Several studies suggested that m^6^A in viral RNA enhances the expression of viral transcripts (50), regulates their splicing (51,52) or promotes interactions between the vRNA and viral polymerase subunits (53). An alternative model postulates that m^6^A regulates the cell-autonomous innate immune response, indirectly facilitating IAV replication by controlling the expression of antiviral proteins (42,54). Thus, whether m^6^A promotes IAV replication primarily through direct effects on viral RNA or indirectly through regulation of host antiviral responses remains unresolved.

In this study, we characterised how m^6^A RNA modifications regulate IAV replication. Inactivation of the METTL3 writer complex or depletion of YTHDF2 decreased viral replication, highlighting that m^6^A RNA modifications are required for viral replication. A quantitative m^6^A mapping technique detected little m^6^A deposition on viral mRNA, though extensive m^6^A modifications were detected on cellular transcripts. Many transcripts encoding antiviral proteins were m^6^A-modified, highlighting how this modification controls the innate immune response. Supporting the role of the m^6^A modifications for controlling cell-autonomous innate immunity, inhibition of type I interferon signalling restored IAV replication when METTL3 activity was inhibited. Analysis of a panel of antiviral proteins that restrict IAV replication revealed that m^6^A inhibition increased expression of the paralogs IFITM2 and IFITM3, and these proteins were required for the antiviral effect of the METTL3 inhibitor. We propose that m^6^A controls IAV replication by acting as a buffer system that limits the expression of a subset of antiviral proteins, including IFITM2/3, thereby balancing the induction of an antiviral response with preventing damage from proteins that alter cellular homeostasis.

## RESULTS

### The m^6^A writer complex and YTHDF2 are required for IAV replication

Previous studies have proposed multiple mechanisms by which m^6^A regulates IAV replication, either by acting directly on viral RNA or indirectly through regulation of the antiviral response (42,50–54). While all proposed mechanisms require the m^6^A writer complex, they differ in their reported dependence on specific reader proteins. For example, m^6^A modifications have been reported to regulate the assembly of the viral ribonucleoprotein (vRNP) complex independently of reader proteins (53). In contrast, IAV pre-mRNA splicing has been linked to the nuclear reader YTHDC1 (51,52), while the cytoplasmic reader YTHDF2 has been implicated in the regulation of both IAV RNA expression and the antiviral response against IAV (42,50,54).

To characterise how perturbations of m^6^A machinery affect IAV replication, we first depleted METTL3 and METTL14 in human A549 lung epithelial cells using siRNAs and infected the cells with IAV (A/WSN/1933(H1N1)) (IAV-WSN) for 48 h at a low multiplicity of infection (MOI = 0.01). This infection condition permits multiple rounds of replication and spread. As expected, METTL3 or METTL14 depletion resulted in a significant reduction in IAV replication (Fig. 1A). We next depleted the canonical m^6^A readers (Fig. 1B). Depletion of YTHDF2 substantially suppressed IAV replication, while depleting YTHDC1 did not significantly reduce viral replication. Therefore, the effect of m^6^A on IAV replication appears to be mediated predominantly through YTHDF2. Of note, targeting the m^6^A eraser ALKBH5 with siRNA did not substantially affect IAV replication (Fig. 1C), consistent with the idea that ALKBH5-mediated demethylation has a minor impact on overall m^6^A levels (20,55).

**Figure 1.**
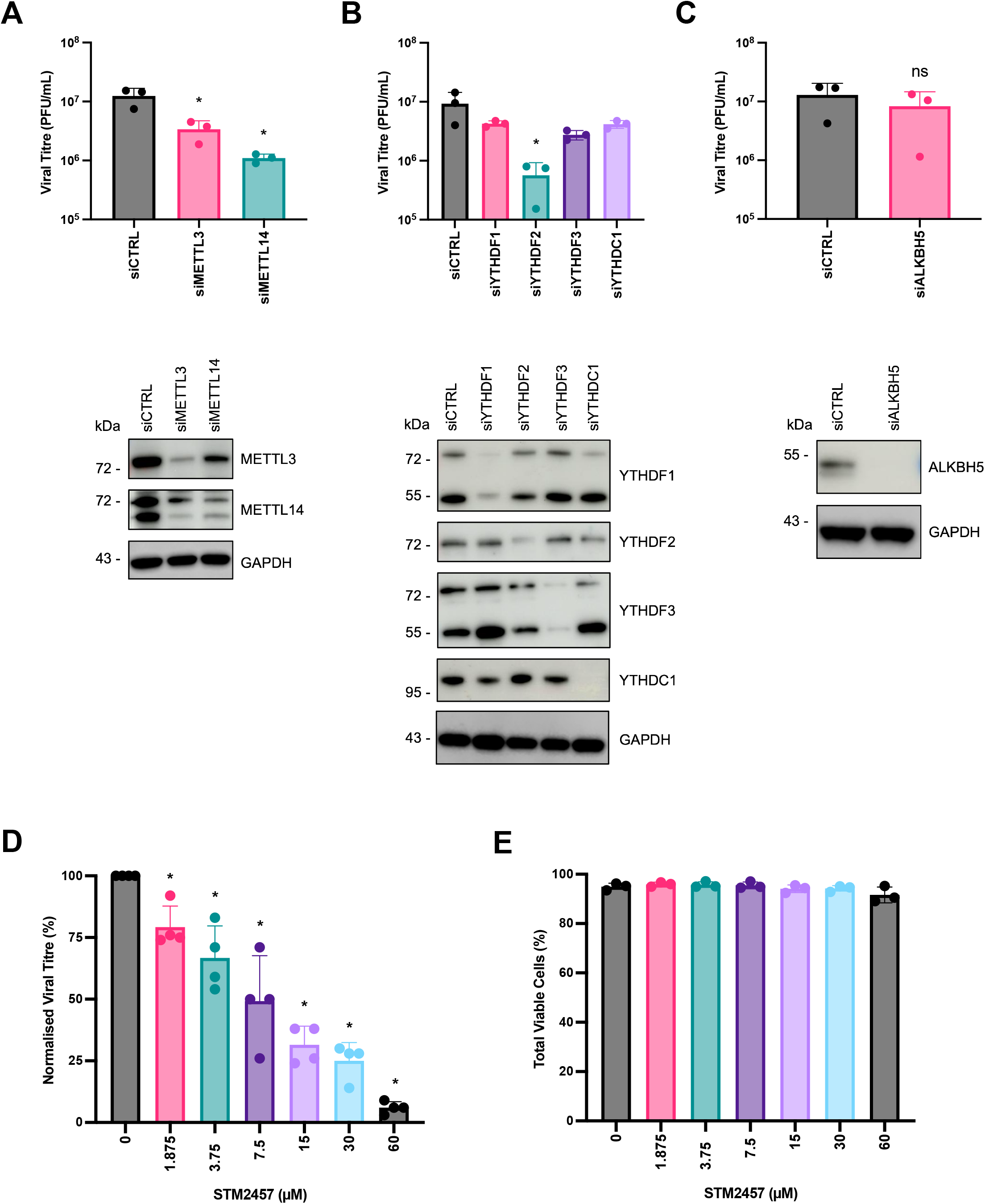
Replication of IAV is modulated by the cellular m^6^A machinery. **(A-C)** A549 cells were transfected twice with siRNAs targeting the indicated components of the m^6^A pathway or a non-targeting control and infected with IAV-WSN at an MOI of 0.01 for 48h. Depletion of the targeted proteins was confirmed by western blotting. **(D)** A549 cells were pretreated for 2 hours with STM2457 at the concentrations ranging from 1.875 μΜ to 60 μΜ or with vehicle control DMSO before infection with IAV-WSN at an MOI of 0.01. Infected cells were maintained in medium containing STM2457 or DMSO for 48h. Viral titres were determined by plaque assay on MDBK cells. Data represent mean ± SD from n = 3 biological replicates. * denotes a statistically significant difference (p < 0.05), as determined by one-way ANOVA of log-transformed data. **(E)** The viability of A549 cells following STM2457 treatment was assessed by flow cytometry using a live/dead cell stain.

While this data supported the notion that m^6^A modifications promote IAV replication, genetic disruption of METTL3 or METTL14 has inherent limitations since most cell lines, including A549, require it for long term survival (56). Consequently, reduced viral replication could partly reflect secondary effects on cellular homeostasis in cells transfected with siRNAs targeting m^6^A writer proteins for several days prior to infection rather than a direct dependence of the virus on m^6^A. We therefore treated A549 cells with the small-molecule inhibitor STM2457 (46), which suppresses METTL3 catalytic activity, and quantified IAV-WSN replication following a 48-hour infection at a low MOI (Fig. 1D). Consistent with our siRNA data, STM2457 treatment significantly restricted IAV replication. Importantly, live/dead cell staining revealed no significant impact of STM2457 on A549 cell viability across all concentrations used for the duration of the assay, indicating that the antiviral effect of the compound is not attributable to general cytotoxicity (Fig. 1E). STM2457 also inhibited IAV (A/California/7/2009(H1N1)) and IAV (A/PR/8/34(H1N1)) (IAV-PR8) (Supplementary Fig. S1A) in A549 cells and IAV-WSN in HBEC3-KT cells, which are a human immortalized, non-cancerous bronchial epithelial cell line (Supplementary Fig. S1B-C), demonstrating that this effect is not strain- or cell type-specific. Together, these findings indicate that the catalytic activity of METTL3 is directly required for efficient IAV replication.

### Influenza A virus transcripts do not contain high stoichiometry m^6^A sites

There are two non-mutually exclusive possibilities for how m^6^A deposition could regulate IAV replication in a YTHDF2-dependent manner: regulation of viral transcripts or regulation of cellular transcripts (42,50,54). While several techniques have been used to identify m^6^A sites on IAV RNA (50,53,57,58), their positions, stoichiometry and importance for viral replication remain unclear. We analysed a previously published set of photo-crosslinking-assisted m^6^A-sequencing (PA-m^6^A-seq) and photoactivatable ribonucleoside-enhanced crosslinking and immunoprecipitation (PAR-CLIP) datasets for ectopically expressed YTHDF1, YTHDF2, YTHDF3 or GFP proteins in cells infected with IAV-PR8 (50) to assess m^6^A modification and reader binding on IAV transcripts. Analysis of the PA-m^6^A-seq dataset indicated that GGACU-containing DRACH motifs were enriched with high PEKA scores (59) (Supplementary Fig. S2A). The PAR-CLIP dataset for YTHDF1 showed similar motif enrichment, though with lower PEKA scores, while the YTHDF2 and YTHDF3 datasets showed weaker enrichment for DRACH sites (Supplementary Fig. 2B-E). This indicates that the PA-m^6^A-seq and YTHDF1 PAR-CLIP datasets are strongly enriched in m^6^A sites, while the PAR-CLIP datasets for YTHDF2 and YTHDF3 show weaker evidence of direct binding at modified sites. We then used *clipplotr* (60) to visualise the PA-m^6^A-seq and PAR-CLIP crosslinks across the IAV cRNA, normalised as a proportion of total crosslinks (Supplementary Fig. S3). Consistent with previous reports (50,53), PA-m^6^A-seq peaks were observed on segments 4, 5 and 6. There was limited overlap with YTHDF1, YTHDF2 or YTHDF3 peaks. However, we noted that the normalized data indicated that even the largest peaks accounted for <0.015% of total crosslinks. As IAV RNA is highly abundant in infected cells (61), it was plausible that only a small proportion of the viral transcripts were m^6^A modified.

To quantitatively analyse m^6^A modifications on IAV mRNAs, we used deamination adjacent to RNA modification targets sequencing (DART-seq) (16,62). In this approach, stable A549 cells expressing a YTH domain-APOBEC1 fusion protein under a doxycycline-inducible promoter were generated and infected with IAV-WSN at a low MOI for 48 hours. The YTH-APOBEC1 fusion protein deaminates cytidine to uridine near m^6^A sites, generating C-to-U substitutions that can be detected by RNA-seq. Importantly, DART-seq is an antibody-independent method for detecting m^6^A sites and therefore provides an orthogonal approach to previous studies of IAV-infected cells using PA-m^6^A-seq or m^6^A immunoprecipitation-sequencing (meRIP–seq) (50,53). Expression of the YTH-APOBEC1 fusion protein did not interfere with IAV replication (Supplementary Fig. S4A-B). Candidate ‘DRACH’ sites were considered m^6^A-modified when C-to-U editing was present in at least 5% of total reads at that site, a threshold determined to effectively eliminate false positives by comparison to conversion rates in cells without induction of YTH-APOBEC1 expression. This threshold is also similar to those used in previous studies (16,62,63). 79,919 unique m^6^A sites were identified in exons across 10,116 genes, with the majority of the modified transcripts harbouring multiple m^6^A sites (Fig. 2A). As expected, these sites displayed characteristic features of m^6^A, including enrichment near the stop codon and within the 3’ UTR (8,9), and were substantially repressed in the presence of STM2457 (Fig. 2B). Cumulative distribution analysis further revealed that most m^6^A sites exhibited relatively low stoichiometry, with highly modified sites representing only a small subset of the transcriptome (Supplementary Fig. S4C). Analysis of the representative cellular transcript ACTB, previously reported to contain m^6^A modifications (10,16,62), showed robust C-to-U editing within its 3’ UTR in both uninfected and infected cells (Fig. 2C-D). In contrast, although low-frequency C-to-U substitutions were detected across IAV transcripts, none exceeded the 5% threshold (Fig. 2E-F), indicating that IAV mRNAs are not substantially modified by m^6^A. Thus, while previous studies have reported the presence of m^6^A on IAV mRNAs (50,57,58), our quantitative DART-seq analysis suggests that modifications occur at very low stoichiometry, which is not consistent with models proposing that m^6^A methylation of viral RNA is required for replication or that it marks viral RNA for decay as part of a surveillance system.

**Figure 2.**
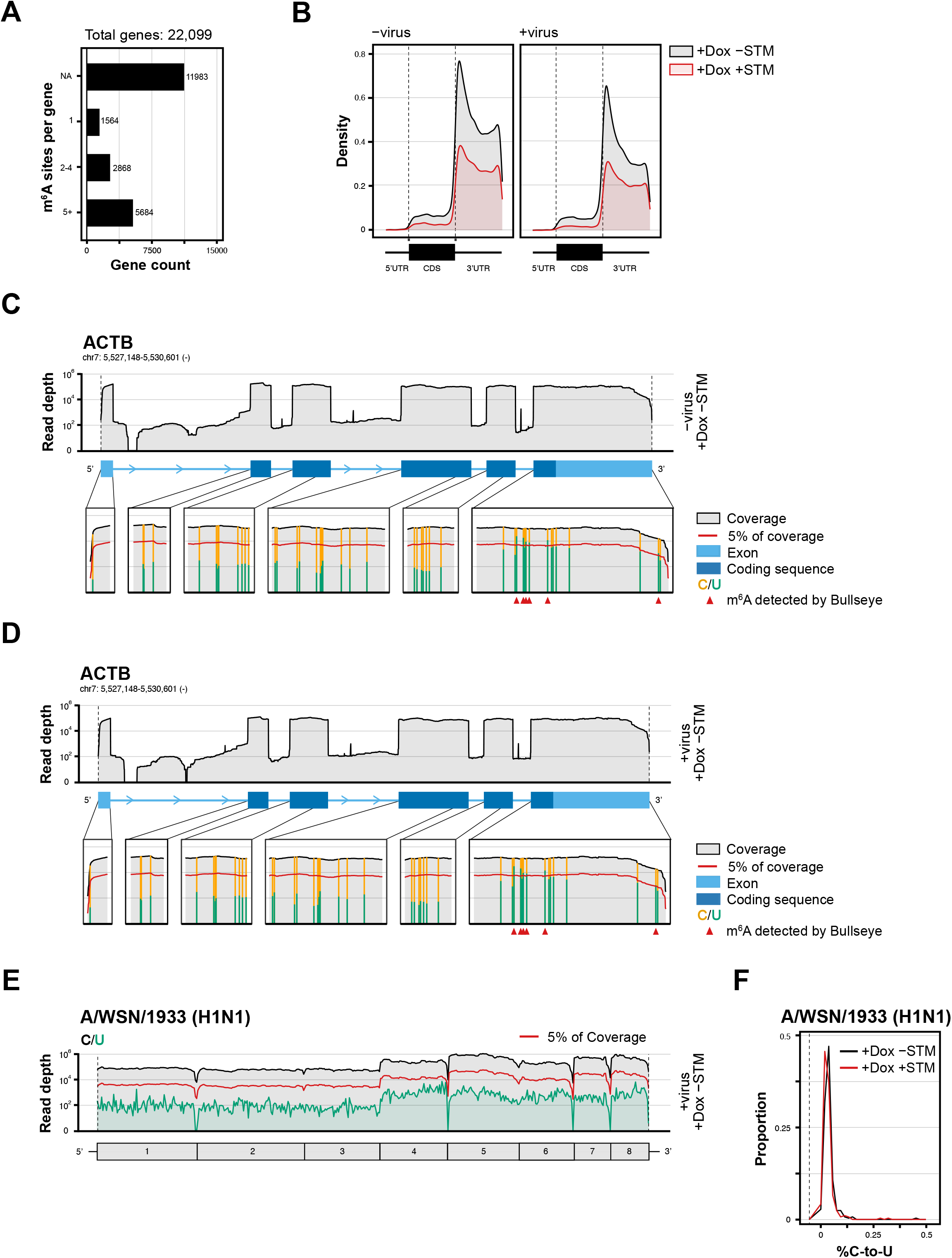
DART-seq reveals widespread m^6^A modification of cellular transcripts but limited modification of IAV mRNAs. DART-seq was performed on poly(A)^+^ RNA extracted from A549 cells expressing a doxycycline (Dox)-inducible YTH-APOBEC1 fusion protein that were either mock- or IAV-WSN-infected (MOI = 0.01, 48 h) in the presence or absence of STM2457 (60 μΜ). **(A)** DART-seq analysis of cellular transcripts. The number of mRNAs containing the indicated number of identified m^6^A sites is shown. Candidate DRACH sites with C-to-U editing in ≥ 5% of reads were classified as m^6^A-modified. **(B)** Metagene analysis depicting the distribution of C-to-U editing events across cellular transcripts in the presence or absence of STM2457. **(C, D)** DART-seq analysis of the representative cellular transcripts ACTB in (C) uninfected and (D) IAV-WSN-infected cells. **(E, F)** DART-seq analysis of IAV-WSN mRNAs showing C-to-U editing frequency in RAC sites across viral transcripts.

### m^6^A regulates the antiviral response to IAV infection

As the viral mRNA was not substantially modified, an alternative explanation for the potent inhibition of IAV replication by METTL3/METTL14 depletion/inactivation or YTHDF2 depletion is that m^6^A regulates cellular transcripts that are either required for viral replication or mediate the antiviral response. To differentiate between these possibilities, A549 cells were infected at a low MOI for 48 hours in the presence of STM2457 alone or in combination with inhibitors targeting antiviral pathways: Ruxolitinib, a JAK inhibitor that blocks type I interferon signalling (64); ISRIB, an integrated stress response inhibitor that counteracts the effects of eIF2α phosphorylation (65); or Bay11-7085, an IκBα phosphorylation inhibitor that disrupts NF-κB signalling (66) (Fig. 3A).

**Figure 3.**
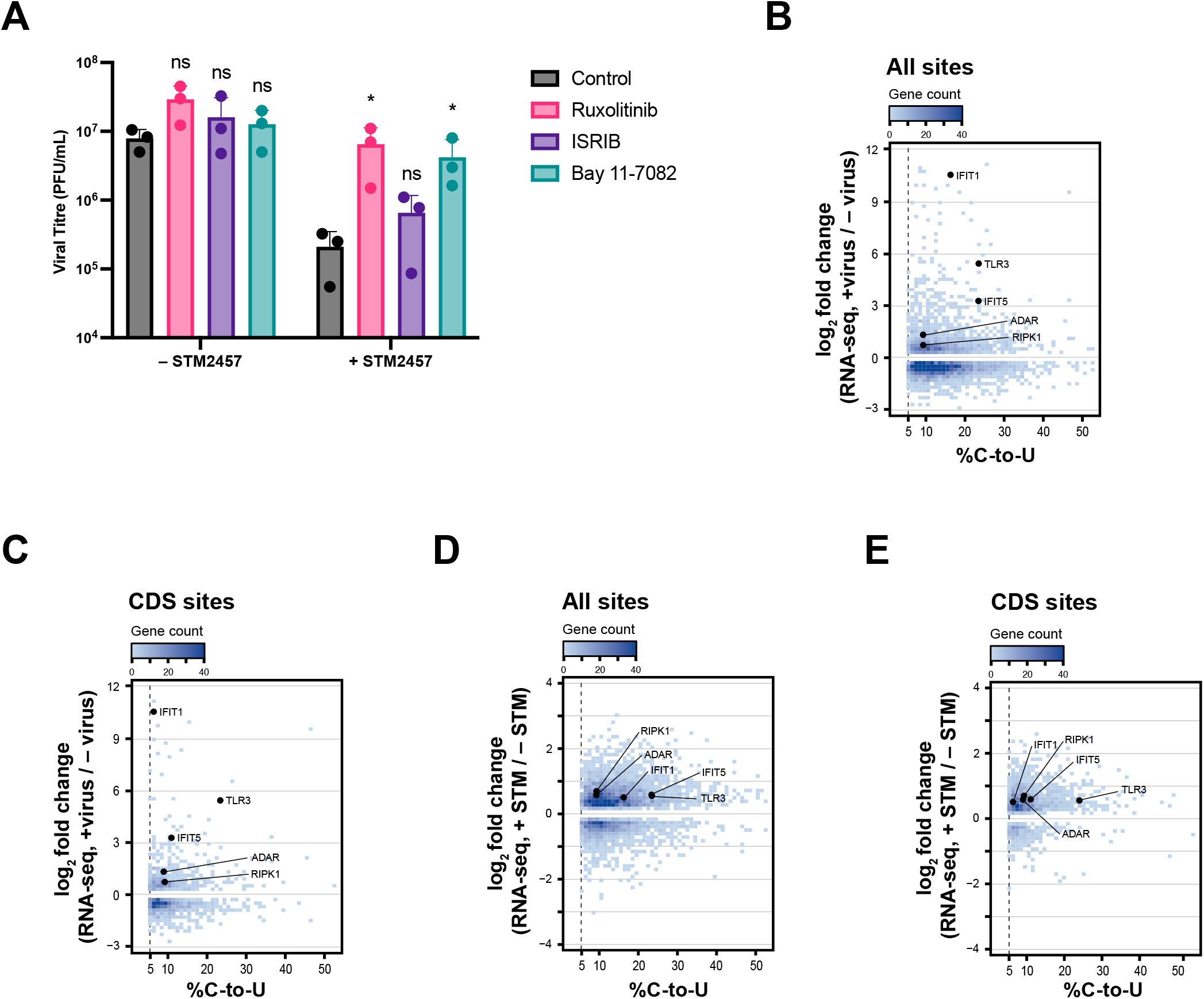
METTL3-14 regulates IAV replication in an interferon-dependent manner. **(A)** A549 cells were pretreated with 60 μΜ STM2457 alone or in combination with 10 μΜ Ruxolitinib, 1 μΜ ISRIB, or 10 μΜ Bay11-7085 for 2 h before infection with IAV-WSN (MOI = 0.01). Infected cells were maintained in medium containing the indicated compounds for 48 h. **(B, C)** Identification of cellular genes upregulated following 48-hour IAV-WSN infection at MOI=0.01 and containing (B) at least one m^6^A site > 5% or (C) at least one m^6^A site > 5% within the coding sequence (CDS). Only genes with adjusted p < 0.05 in the + virus / - virus RNA-seq comparison are shown. Genes encoding proteins that regulate programmed necrotic cell death (RIPK1), modulate viral replication (ADAR, IFIT1, IFIT5), or encode a pattern recognition receptor (TLR3) are highlighted. **(D, E)** Effect of STM2457 treatment on the expression of genes identified in Fig. 3B-C.

Strikingly, treatment with Ruxolitinib potently rescued IAV replication in STM2457-treated cells, suggesting that the restriction of IAV in m^6^A-deficient cells requires type I interferon signalling. Bay11-7085 partially rescued IAV from the METTL3 inhibition, while ISRIB had no significant effect on viral propagation (Fig. 3A).

Since Ruxolitinib and Bay11-7085 increased viral replication in STM2457-treated cells, we hypothesized that m^6^A modulates the antiviral response to IAV infection. Analysis of our DART-seq dataset supported this idea (Fig. 3B-C). Of the 5,886 virus-induced genes (VIGs, log_2_ fold change > 0.5, padj < 0.05), 1,094 had at least one m^6^A site in an exon (editing rate > 5%) (Fig 3B). As m^6^A sites in the coding sequence (CDS) have been reported to mediate RNA decay more effectively than those within the 3’ UTR due to co-translational degradation mediated by YTHDF proteins (31,32,67), we also identified 401 VIGs containing m^6^A sites within their CDS (Fig. 3C). Therefore, ∼20% of VIGs had at least one m6A site in an exon and 7% contain at least one site in the CDS. This indicates that m^6^A has the potential to modulate the cellular response to IAV infection. Gene ontology analysis revealed that antiviral terms are strongly enriched amongst the m^6^A-modified VIGs (Supplementary Table S1), including programmed necrotic cell death (FAS, RIPK1), regulation of RIG-I signalling pathway (BIRC2, BIRC3, DHX58, USP15) and regulation of viral genome replication (ADAR, IFIT1, IFIT5, N4BP1, OAS1, OAS2, RNASEL). Genes encoding other proteins that restrict viral replication were also identified (TLR3, SAMD9, SAMD9L, PARP14). Notably, several of these were upregulated upon STM2457 treatment of IAV-infected cells, including RIPK1, ADAR, IFIT1, IFIT5, and TLR3 (Fig. 3D-E), which contain multiple m^6^A sites (Supplementary Fig. S5).

### METTL3 inhibition does not enhance interferon-β production following IAV infection

As m^6^A modifications were detected on mRNAs encoding proteins involved in virus sensing through PRRs as well as proteins that directly inhibit specific steps of viral replication, altered expression of either group could contribute to the antiviral effect of m^6^A inhibition. To determine whether sensing of IAV infection was modulated by m^6^A modifications, we infected cells with IAV-WSN at a high MOI (MOI = 5) and measured interferon beta 1 (IFNB1) mRNA expression 8 hours post-infection, thereby minimising the confounding effects of cumulative rounds of viral replication. STM2457 treatment had no effect on infectious virus production under these conditions (Fig. 4A), indicating that the antiviral effects of METTL3 inhibition emerge predominantly during subsequent rounds of infection and spread (Fig. 1D and 3A). As expected, IAV infection potently induced IFNB1 mRNA expression, but there was no significant difference in the magnitude of induction when m^6^A writing was inhibited (Fig. 4B). Although IFNB1 mRNA has previously been proposed to be m^6^A-modified (42,43), we did not detect any sites with an editing frequency >5% in our DART-seq data (Fig. 4C), nor were m^6^A sites detected on IFNB1 mRNA in the published PA-m^6^A-seq dataset (50) (Supplementary Fig. S6). In contrast to the high MOI experiment (Fig. 4B), IFNB1 induction was moderately decreased in STM2457-treated cells 48 hours after a low MOI IAV-WSN infection (Fig. 4D). Similarly, reduced IFNB1 mRNA expression was also observed in our DART-seq dataset at 48 h post-infection (Fig. 4E). Since we did not find evidence of high frequency m^6^A sites on IFNB1 mRNA, we propose that the reduction in IFNB1 expression in the low MOI experiments following STM2457 treatment is due to reduced viral replication, leading to less PAMP being expressed, rather than direct effects of m^6^A on IFNB1 mRNA stability. These results demonstrate that inhibition of m^6^A writer activity does not substantially alter detection of IAV infection.

**Figure 4.**
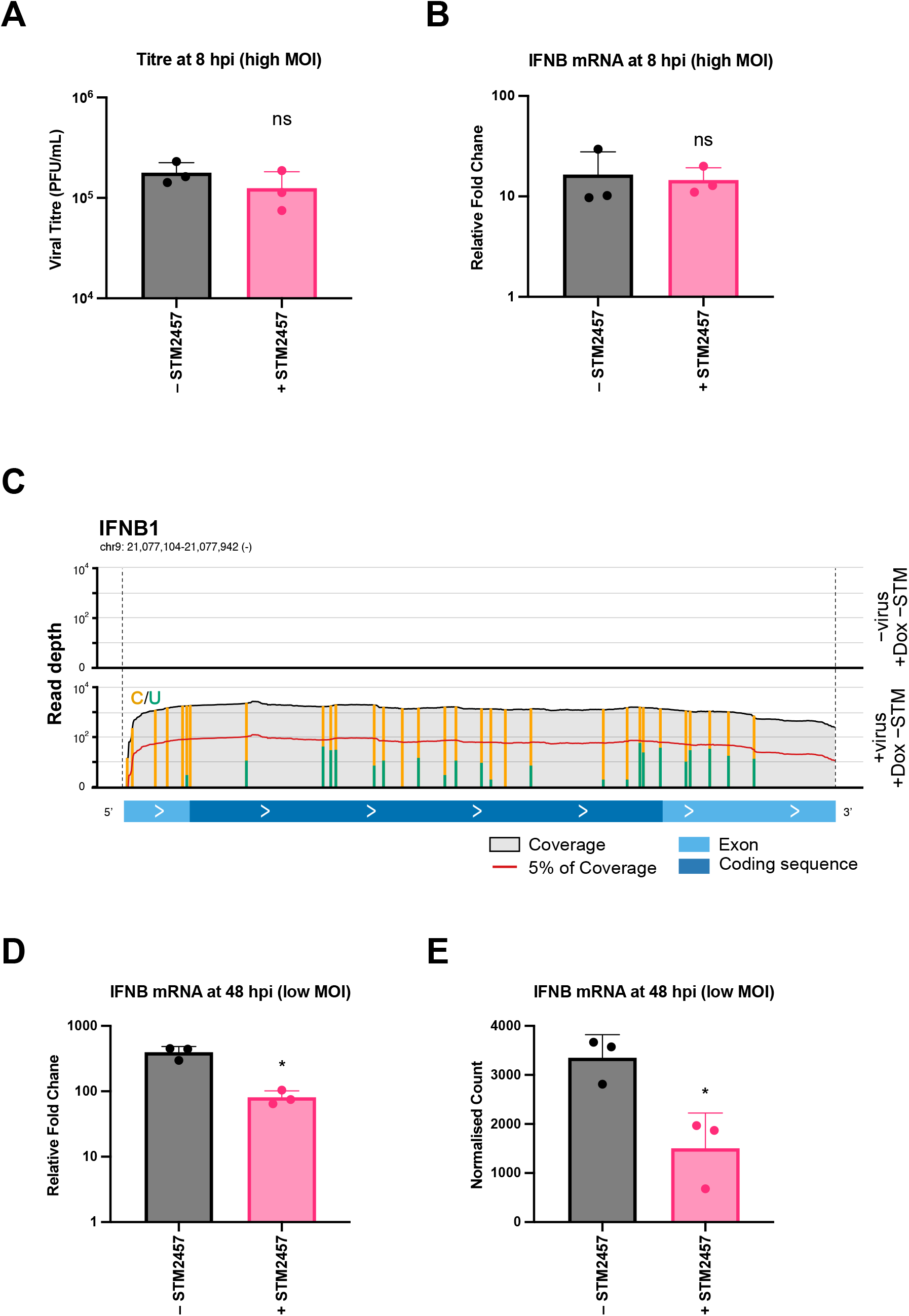
METTL3-14 inhibition does not enhance interferon induction following IAV infection. **(A-B)** A549 cells pretreated for 2 hours with STM2457 (60 μΜ) or DMSO before infection with IAV-WSN at an MOI = 5. Infected cells were maintained in the presence of STM2457 or DMSO for 8 hours. (A) Viral titres were determined by plaque assay on MDBK cells. (B) IFNB1 mRNA induction was measured by RT-qPCR and normalised to 18s rRNA. **(C)** DART-seq analysis of the IFNB1 transcript. **(D-E)** IFNB1 mRNA abundance 48 hours post-infection (hpi) with IAV WSN (MOI = 0.01) in the presence or absence of STM2457, as determined by (D) RT-qPCR, normalised to 18S rRNA, and (E) RNA-seq.

Although IFNB1 mRNA levels were decreased in STM2457-treated cells at 48h post-infection, it remained strongly expressed. Consistent with this, western blotting showed that the PRRs MDA5 and RIG-I, which are also ISGs, were robustly induced in both control and STM2457-treated cells at 48 h post-infection (Fig. 5A). No m^6^A sites were detected in RIG-I mRNA, while those identified in MDA5 mRNA were located outside the CDS (Supplementary Table S1). As expected, MAVS expression was decreased by IAV infection, potentially via PB1-F2 (68), and this decrease was less evident with STM2457 treatment, consistent with lower viral replication. Collectively, these findings indicate that IAV attenuation in m^6^A-deficient cells is not driven by enhanced interferon expression or PRR induction.

**Figure 5.**
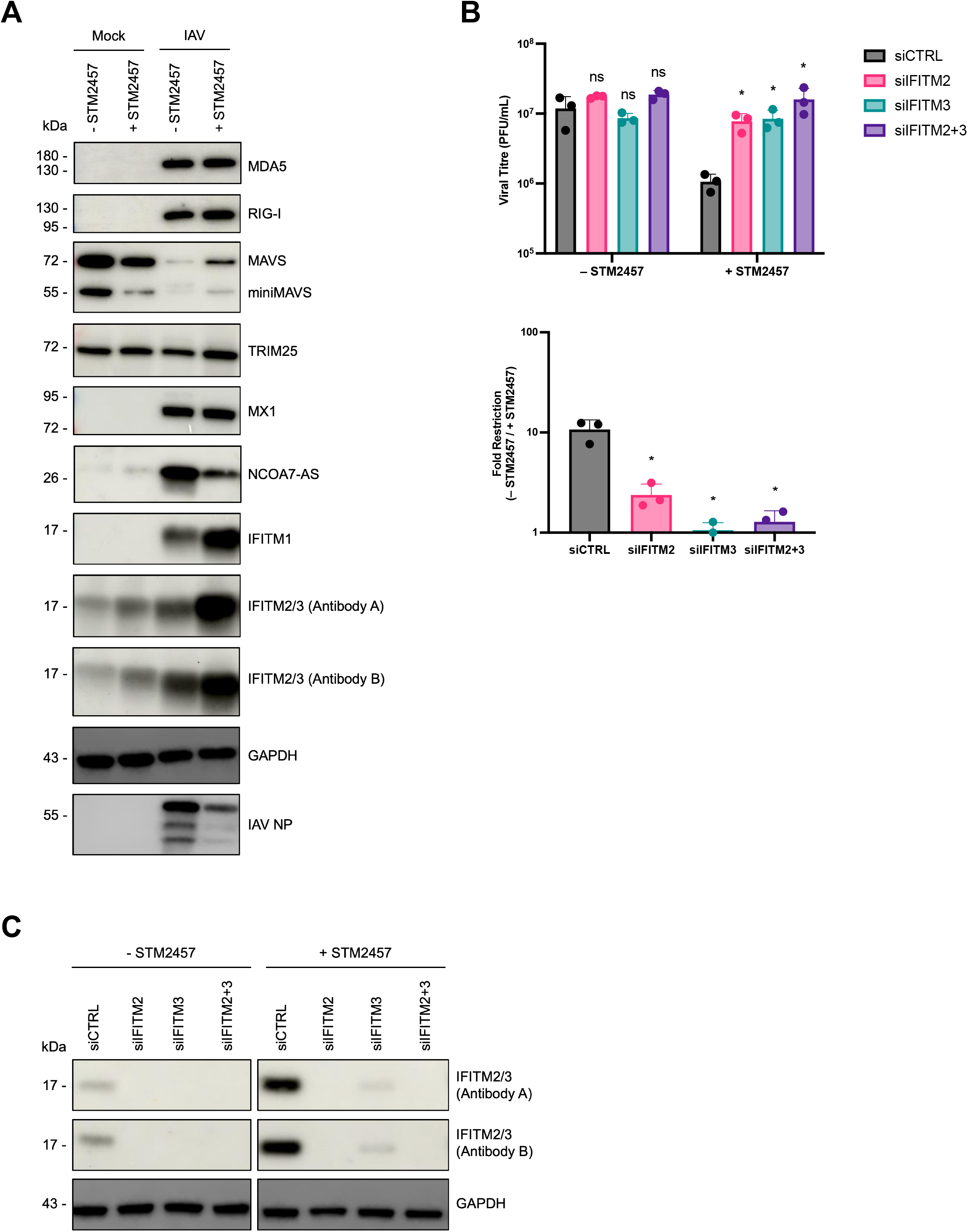
IFITM2/3 mediate IAV restriction following METTL3-14 inhibition. **(A)** Western blot analysis of the indicated cellular proteins in mock- or IAV-WSN-infected A549 cells treated with STM2457 (60 μΜ) or DMSO. Cells were collected 48 hpi. Antibody A is 12769-I-AP, antibody B is 11714-I-AP. **(B, C)** A549 cells were transfected twice with siRNAs targeting IFITM2, IFITM3 or both, or with a non-targeting control siRNA, and subsequently pretreated for 2 hours with 60 μΜ STM2457 or DMSO before infection with IAV-WSN at an MOI of 0.01. Infected cells were maintained in the presence of the drug for 48 hours. (B) Viral titres were determined by plaque assay on MDBK cells (top panel) and STM2457-mediated restriction of viral replication (bottom panel) are shown. (C) IFITM2/3 knockdown efficiency assessed by western blot. Antibody A is 12769-I-AP, antibody B is 11714-I-AP.

### METTL3 inhibition increases IFITM2/3 expression

Since the inhibitory effect of STM2457 did not appear to be explained by increased interferon expression, we hypothesized that ISGs that directly target IAV replication might be affected. Similar to RIG-I and MDA5, induction of MX1, which restricts IAV by targeting vRNPs (69,70) was unaffected by STM2457 (Fig. 5A), further supporting the observation that altered interferon expression is not responsible for the effect of m^6^A inhibition on viral replication. NCOA7-AS, the interferon-inducible isoform of NCOA7 that restricts IAV entry via endosomes (71), was moderately reduced by STM2457 treatment (Fig. 5A), making it unlikely to account for the antiviral phenotype. Interestingly, treatment with the METTL3 inhibitor increased IFITM1, IFITM2 and IFITM3 (Fig. 5A). IFITM2 and IFITM3 are highly related interferon-induced endosomal transmembrane proteins that inhibit IAV cell entry by preventing fusion between the viral and endosomal membranes (72–75). Although IFITM2 and IFITM3 are two distinct proteins, their high sequence homology results in antibody cross-reactivity, and the proteins are therefore referred to collectively as IFITM2/3 where they cannot be distinguished. IFITM2/3 expression was upregulated by STM2457 in both uninfected and IAV-infected cells (Fig. 5A). To determine whether IFITM2/3 were required for STM2457-mediated restriction of IAV, we knocked them down individually or in combination using siRNAs. Remarkably, IFITM2/3 knockdown fully restored viral replication in STM2457-treated cells (Fig. 5B-C). This indicates that the reduction of IAV replication when m^6^A writing is inhibited is mediated predominantly through upregulation of IFITM2/3, revealing that m^6^A modifications regulate the expression of antiviral proteins.

We then analysed IFITM2 and IFITM3 transcript levels. We had previously observed that m^6^A writer inhibition did not restrict IAV replication in high MOI experiments (Fig. 4A), while the virus was strongly restricted in low MOI experiments where there are multiple rounds of viral replication (Fig. 1D and 3A). Consistent with this, IFITM2 and IFITM3 were not induced at 8 hours in a high MOI experiment (Fig. 6A), likely due to a temporal lag between interferon expression and ISG induction. However, they were potently induced at 48 hours post-infection in a low MOI experiment (Fig. 6B). Interestingly, STM2457 treatment did not alter IFITM2 or IFITM3 mRNA induction under either experimental condition, suggesting that the increase in IFITM2/3 expression is not due to increased transcript abundance. Analysis of the DART-seq dataset showed that there were no m^6^A sites in IFITM2 or IFITM3 above the 5% editing threshold (Fig. 6C-D), indicating that they are not directly modified by METTL3. Similarly, no m^6^A sites were detected on IFITM2 or IFITM3 transcripts in the published PA-m^6^A-seq dataset (50) (Supplementary Fig. S7A-B). Since METTL3-14 inhibition increased IFITM2/3 protein levels without affecting their mRNA abundance, we analysed their protein half-life using a cycloheximide chase assay. We found that STM2457 treatment significantly extended the half-life of IFITM2/3 (Fig. 6E, Supplemental Fig. 8A), whereas no substantial change in protein half-life was observed for MX1 (Fig. 6F, Supplemental Fig. 8A).

**Figure 6.**
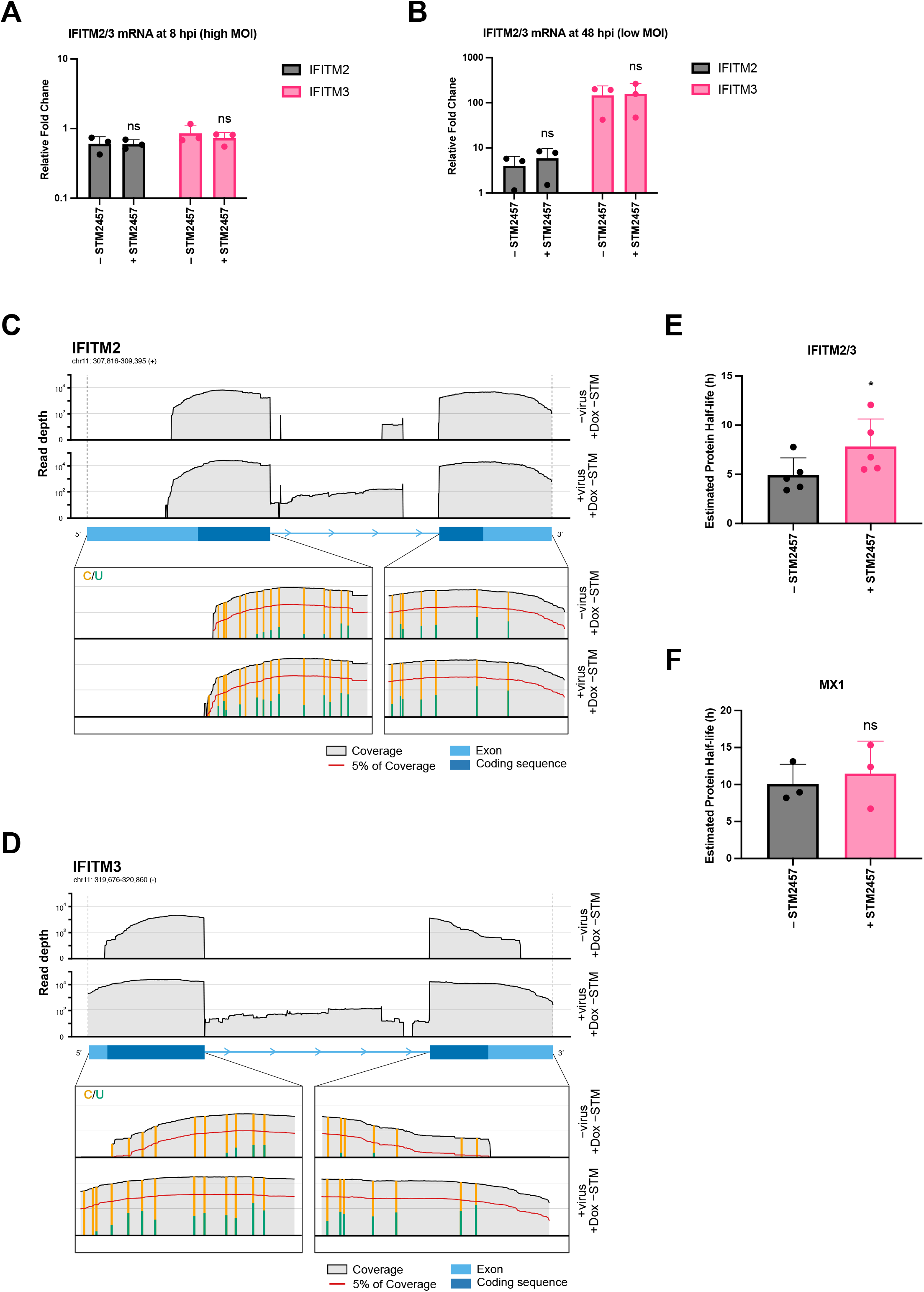
METTL3-14 inhibition increases IFITM2/3 protein stability. **(A, B)** IFITM2 and IFITM3 mRNA induction in A549 cells infected with IAV-WSN in the presence or absence of 60 μΜ STM2457 under (A) high-MOI (MOI = 5, 8 hours) or (B) low-MOI (MOI = 0.01, 48 hours) conditions, as determined by RT-qPCR. **(C, D)** DART-seq analysis of (C) IFITM2 and (D) IFITM3 transcripts. **(E, F)** Protein half-life of (E) IFITM3/2 and (F) MX1 in A549 cells pretreated with IFN-β (50 U/mL) and STM2457 (60 μΜ) or DMSO, as determined by cycloheximide chase followed by western blotting.

The mechanisms regulating IFITM2/3 protein stability remain poorly understood. Ubiquitylation by NEDD4 has been reported to regulate IFITM3 abundance and S-palmitoylation increases IFITM3 antiviral activity (76–78). However, NEDD4 depletion did not affect STM2457-mediated restriction of IAV replication and pharmacological inhibition of palmitoylation was toxic in IAV-infected A549 cells (Supplemental Fig. S8B-C). Interestingly, cyclosporine H (CsH) or its derivatives have been shown to antagonise IFITM3-mediated restriction of lentivector transduction through unknown mechanisms (79,80). While CsH itself has direct antiviral properties against IAV (81), it abrogated the antiviral effect of STM2457 (Supplemental Fig. S9A-B), further supporting our conclusion that METTL3 inhibition restricts IAV replication through regulation of IFITM2/3 expression.

Overall, we propose a model in which m^6^A promotes IAV replication by limiting the accumulation of the antiviral proteins IFITM2/3 (Fig. 7). During the initial exposure to the virus (first round of infection), cells express low levels of IFITM2/3 and are therefore permissive to viral replication. However, as infection progresses and type I interferon signalling induces IFITM2/3 expression, METTL3 inhibition enhances IFITM2/3 protein stability, leading to their accumulation in infected cells. Elevated IFITM2/3 levels subsequently restrict viral entry during secondary rounds of infection, thereby limiting viral spread.

**Figure 7.**
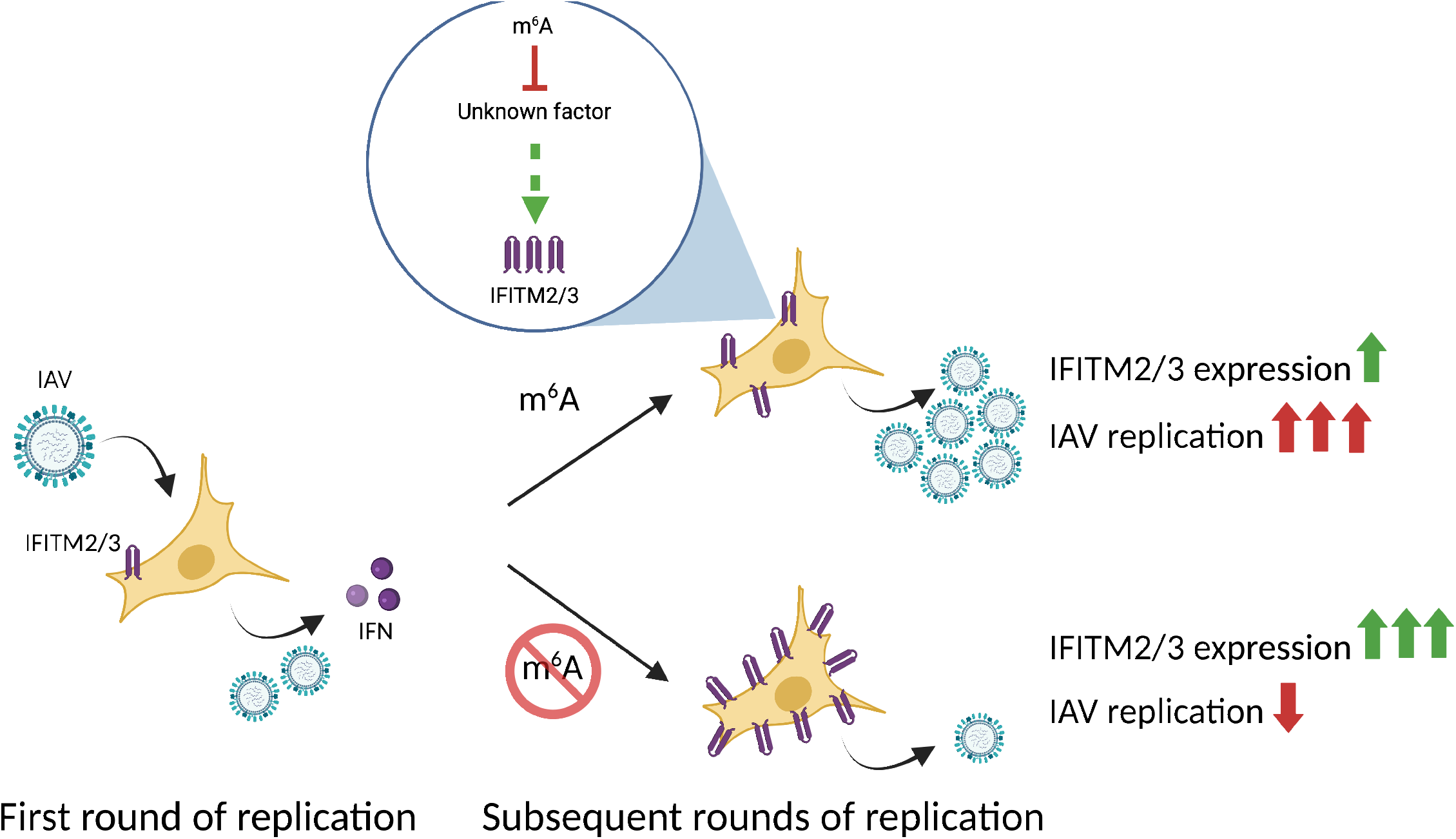
Proposed model of m^6^A-mediated regulation of IAV replication through IFITM2/3.

## DISCUSSION

There are several proposed roles for m^6^A-mediated control of gene expression, including regulating cellular differentiation and surveillance of improperly processed transcripts or viral RNA (22,33). In this study, we investigated how m^6^A controls IAV infection. While it has been proposed that the viral RNA is directly modified (50–53), m^6^A stoichiometry had not been previously measured at any of the proposed sites. Using DART-seq, we did not identify any m^6^A sites in IAV mRNA with a frequency > 5%. We note that this analysis did not include IAV vRNA or cRNA, which are not polyadenylated, though we did not identify any high abundance peaks when we reanalysed PA-m^6^A-seq and YTHDF1, YTHDF2 or YTHDF3 PAR-CLIP datasets in IAV infected cells (50). IAV vRNA and cRNA are co-transcriptionally coated with the viral nucleoprotein, and it is unclear how the m^6^A writer complex would efficiently access and modify these transcripts in the context of the known mechanisms for viral genome replication (82,83). Overall, the low rate of m^6^A modification of IAV RNA is not consistent with it directly promoting specific steps of viral replication or targeting these transcripts for decay as part of an RNA surveillance system. While it was originally proposed that m^6^A modifications are a common feature for RNA virus transcripts based on experiments using antibody-based methods to identify m^6^A sites (34), this has been re-evaluated using antibody-independent technologies for some positive strand RNA viruses and viral RNAs may be considerably less modified than originally believed (84). Comparing our DART-seq data for IAV-infected cells to previous reports using PA-m^6^A-seq or meRIP–seq (50,53) is consistent with the observation that antibody-based m^6^A mapping approaches may provide an overestimation of m^6^A sites in viral RNA (84), which are highly abundant transcripts in infected cells, highlighting the importance of using orthogonal approaches.

Our data indicate that m^6^A-mediated modulation of IFITM2/3 expression is responsible for the inhibition of IAV replication observed when the m^6^A writer complex is inhibited (42,50–54). Importantly, depletion of IFITM2/3 completely rescued IAV replication in STM2457-treated cells, suggesting that increased IFITM2/3 abundance is sufficient to account for the antiviral effect of METTL3 inhibition under these conditions. This does not exclude contributions from previously proposed mechanisms (42,50–54) under other experimental conditions but indicates they are not required to explain the antiviral phenotype observed here. IFITM proteins inhibit a wide range of viruses that enter the cell via membrane fusion (73,75). IFITM2/3 restrict IAV replication, and polymorphisms in IFITM3 are linked to IAV morbidity from emerging IAV strains (85,86). Regulating IFITM2/3 expression levels may be important for cellular homeostasis because these proteins disrupt local lipid sorting to inhibit the fusion of viral and cellular membranes (74), but their excessive or prolonged expression may interfere with essential cellular processes, such as membrane trafficking or cell-cell fusion. For instance, elevated IFITM expression has been reported to impair cell-cell fusion events required for normal placenta development, contributing to placental pathology associated with congenital viral infections or aberrant interferon responses (87,88). In this context, mechanisms that restrain IFITM2/3 abundance following interferon-driven induction may be important for balancing antiviral protection with the need to preserve cellular homeostasis. How m^6^A modifications control IFITM2/3 expression remains unclear. m^6^A did not appear to directly modulate the IFITM2 or IFITM3 mRNA abundance, and there were no detectable m^6^A modification sites > 5% on either transcript. Therefore, we hypothesize an IFITM2/3 cofactor that regulates its half-life may be directly regulated by the m^6^A pathway. Substantial knowledge gaps remain for how IFITM2/3 expression is regulated, including how cyclosporin H targets IFITM2/3 for degradation (79). As the STM2457-mediated restriction of IAV replication is lost after cyclosporin H treatment, the transcript encoding this unknown factor may be m^6^A-modified. Identifying the m^6^A-regulated factor(s) responsible for controlling IFITM2/3 protein stability will therefore be an important direction for future investigation.

Control of cell-autonomous innate immune responses by m^6^A modifications is important for a wide variety of conditions ranging from viral infection to type 1 diabetes (36,38,89). Our DART-seq analysis extends these findings by providing quantitative information on m^6^A modification across virus-induced cellular transcripts. As m^6^A sites in the CDS have a more prominent effect on RNA decay than those in the 3’ UTR (31,32,67), we stratified our data and found that approximately 7% of VIGs have at least one CDS site with >5% modification. This includes transcripts encoding known proteins that mediate or regulate the antiviral response, including ADAR, IFIT1, IFIT5, RIPK1 and TLR3, consistent with previous findings (38,89). We propose that m^6^A-mediated RNA decay buffers the cell-autonomous innate immune response for a subset of proteins that sense or restrict viral replication by limiting the amount of protein produced per transcript. m^6^A sites in these mRNAs will increase the probability of ribosome stalling and collisions, leading to YTHDF-mediated RNA decay (31,32). This may be an overall role for m^6^A-mediated post-transcriptional gene regulation in that it could buffer protein expression when specific transcriptional programmes are induced, including those in cell differentiation and stress responses (22,32). Other transcripts encoding antiviral proteins contained m^6^A sites predominantly in their 3’ UTR, and the relative functional contribution of these modified residues requires further investigation. However, m^6^A sites in the 3’ UTR may cluster with other post-transcriptional regulatory elements, such as AU-rich elements, which can lead to highly unstable transcripts under specific biological conditions (90). Further investigation of how cell-autonomous innate immunity is controlled by m^6^A alone and in combination with other post-transcriptional regulatory pathways will help elucidate how the innate immune response is regulated to mediate an efficient anti-pathogen response without causing undue tissue damage.

## METHODS

### Cells

A549 cells were obtained from the ATCC. MDBK and MDCK cells were a kind gift from Hannah Mischo and Wendy Barclay, respectively. They and their derivative cell lines were cultured in DMEM + GlutaMAX (Gibco) supplemented with 10% heat-inactivated foetal bovine serum (Gibco) plus 100 U/mL penicillin and 100 µg/mL streptomycin (Pen Strep, Gibco). HBEC3-KT cells were a gift from Rocio Martinez-Nunez and were grown on collagen-coated plates (Collagen I bovine, Gibco) in AEGM with supplement mix (PromoCell). All cell lines were maintained at 37°C in 5% CO_2_ and routinely tested for mycoplasma infection using PCR Detection Kit (Thermo Scientific Chemicals, #J66117).

Because the A549 cells were also used for SARS-CoV-2 experiments not described in this manuscript, they were engineered to express ACE2 and TMPRSS2 by lentiviral co-transduction with pCMS28-ACE2 and RRL.sin.cPPT.SFFV/TMPRSS2.IRES-neo.WPRE (a gift from Dr. Caroline Goujon, University of Montpellier). The A549-ACE-TMPRSS2-YTH-APOBEC1 cell line used for DART-seq was subsequently generated by transducing A549-ACE2-TMPRSS2 cells with TLCV2-APOBEC1-YTH (Addgene, #178949). For each construct, lentiviral particles were generated using a gag-pol packaging plasmid psPAX2 (Addgene, #12260) and VSV-G envelope plasmid pMD2.G (Addgene, #12259). Transduced cells were subsequently selected using the appropriate antibiotics.

### IAV infection

A full-length influenza A/WSN/1933 (H1N1) was produced using the 12-plasmid-based rescue system (91). Influenza A/Puerto Rico/8/1934 (H1N1) and A/California/07/2009 (H1N1) strains were kindly provided by Hannah Mischo. A549 or HBEC3-KT cells were seeded in a 24-well plate at a density of 8x10^4^ or 3x10^4^ cells per well, respectively. 24 hours after seeding, cells were washed with PBS (Gibco) and infected with IAV diluted in serum-free DMEM to achieve an appropriate MOI (indicated in the relevant figure legends) for 1h at 37°C. The inoculum was then removed and 500 µL of fresh Opti-MEM + GlutaMAX (for A549s) or BEGM (for HBEC-3KTs) was added. For infections with PR8 or CA09 strains, the medium was additionally supplemented with 0.6 µg/mL of TPCK-treated trypsin (Worthington Biochemical). Plates were incubated for 8 hours or 48 hours at 37°C, after which both cells and supernatants were collected and stored at -80°C for downstream analysis.

### RNA interference

4x10^4^ of A549 cells were reverse-transfected with 20 pmol of siRNA (listed in Table S2) and 1 µL of Lipofectamine RNAiMAX (Invitrogen) in a 24-well plate. After 24 h, the cells were subjected to a forward transfection using the same amount of siRNA and Lipofectamine RNAiMAX. 24h later, cells were infected with IAV as described above.

### Small molecular inhibitor treatments

For infection experiments involving STM2457 (Medchem Express, #HY-134836), Ruxolitinib (Selleck Chemicals, #S1378), ISRIB (Sigma, #SML0843), Bay 11-7083 (Cayman Chemical, #CAY14795), PP2 (Abcam, #AB120308) and 2-BP (Medchem Express, #HY-111770) cells were pre-treated with the respective compound(s) for 2 h prior to infection at the concentrations indicated in the figure legends. In the case of Cyclosporine H (Sigma, #SML1575), cells were pre-treated with the compound overnight. DMSO was used as a vehicle control at the corresponding final concentration (% v/v). Following pre-treatment, cells were inoculated with the virus and subsequently maintained in medium containing the same amount of the drug throughout the course of infection.

### Plaque assay

MDBK (for WSN) or MDCK (for PR8 and CA09) cells were seeded in 6-well plates and allowed to grow to 90-95% confluence. Approximately 24 hours later, the cell monolayers were washed with PBS and infected with 400 µL of 10-fold serial dilutions of virus supernatants in serum-free DMEM. The plates were incubated for 1 hour at 37°C, with rocking every 15 minutes to ensure even distribution of the inoculum. Following the incubation period, the infectious media was removed, and 2 mL of the solid overlay medium was gently applied over infected cells. The overlay medium was composed of 1x Minimum Essential Medium (MEM, Gibco) supplemented with 0.21% bovine serum albumin (BSA fraction V, Gibco), 2.0 mM L-glutamine (Gibco), 0.15% sodium bicarbonate (Gibco), 10 mM HEPES (Gibco), 0.005% dextran (Sigma, cat. #D9885), 1x Pen Strep (Gibco), and 0.6% agarose (Ultrapure, Invitrogen). The plates were incubated for 72 hours at 37°C, after which the overlay was carefully removed, and cells were fixed with 0.1% crystal violet solution dissolved in a 20% methanol solution for 15 min at room temperature.

### Viability assay

Cells were seeded in the morning in a 24-well plate at a density of 8x10^4^ (A549) or 3x10^4^ (HBEC3-KT) cells per well. 8 hours later, cells were treated with STM2457 or DMSO control. After a 48-hour incubation at 37°C, the culture medium was collected, and the cells were detached with TrypLE Express (Gibco) directly into the same collection tube. The cells were then stained with the LIVE/DEAD fixable near-IR fluorescent reactive dye (Life Technologies, #L10119), according to the manufacturer’s protocol. Finally, cells were transferred to FACS test tube (Fisher Scientific) and the fluorescence measured using BD FACSCanto II flow cytometer (BD Biosciences). The data were processed using FlowJo (FlowJo LLC).

### Western blot

Cells were lysed in 2x Laemmli sample buffer and denatured for 5 min at 95°C. Protein samples were separated on mPAGE 8-16% Bis-Tris precast polyacrylamide gel (Millipore) and transferred onto either a 0.45 µm PVDF membrane (Cytiva) or 0.45 µm nitrocellulose membrane (Cytiva) using a Criterion Blotter tank transfer system (Bio-Rad). Membranes were blocked for 1 hour at room temperature in 5% skimmed milk (Marvel) prepared in PBST (PBS containing 0.1% Tween 20), followed by incubation with primary antibodies overnight at 4°C and secondary antibodies for 1 hour at room temperature. Membranes were washed four times for 5 minutes in PBST after each antibody incubation. Membranes incubated with IRDye-conjugated secondary antibodies were imaged using an Odyssey Infrared Imagining System (LI-COR Biosciences). For chemiluminescence detection, membranes probed with HRP-conjugated secondary antibodies were treated with Amersham ECL Prime Western Blotting Detection reagent (Cytiva, #RPN2232) for 5 min and imaged using the Amersham Image Quant 800 (Cytiva). Antibodies used in this study are listed in Table S3.

### RNA extractions

Cells were lysed in RLT lysis buffer (Qiagen) supplemented with 10 µL/mL of β-mercaptoethanol. Total RNA isolation was performed using RNeasy Mini Kit (Qiagen) according to the manufacturer’s instructions. RNA was eluted in nuclease-free water and its concentration and integrity assessed using Qubit RNA HS Assay Kit (ThermoFisher, Q32852) and RNA ScreenTape assay (Agilent), respectively.

### RT-qPCR

1 µg of extracted RNA was reverse-transcribed into cDNA with the High Capacity cDNA Reverse Transcription Kit (Applied Biosystems) with random primers, according to the manufacturer’s instructions. The resulting cDNA was then diluted 1:20 in nuclease-free water and used as a template for qPCR reactions. Reactions were performed using TaqMan Gene Expression Assay (Applied Biosystems) with commercially available probes (listed in Table S4). Amplification was carried out using a QuantStudio 5 Real-Time PCR System (Applied Biosystems). Gene expression data were analysed using the Thermo Fisher Connect Platform, and relative expression was determined using the comparative Ct (ΔΔCt) method.

### Cycloheximide chase

A549 cells were seeded in a 24-well plate at of 1x10^5^ cells per well and pre-treated with 50 U/mL of IFN-β (PBL Assay Science, #11415-1) in the presence of 60 µM STM2457 or DMSO control. The following day, cells were treated with 50 µg/mL cycloheximide (CHX). CHX was added to individual wells at 2-hour intervals, with all samples collected simultaneously 10h after initiation of the first CHX treatment. Cells were lysed directly in 2x Laemmli sample buffer, and protein abundance was analysed by western blotting as described above. Band intensities were quantified using Image Studio Lite (version 5.2.5, LI-COR Biosciences). IFITM abundance was normalised to GAPDH and expressed relative to the 0 hour time point for each condition. Protein half-lives were determined by fitting a one-phase exponential decay model to the normalised values in GraphPad Prism (version 10.4.1, GraphPad Software).

### DART-seq analysis

A modified reference genome was constructed by appending the IAV WSN genome (GCA_000001405.29) to the human reference genome (GRCh38.p14). A corresponding modified annotation file was generated by appending viral transcript annotations to the GENCODE v46 human annotation.

Raw reads were trimmed using TrimGalore v0.6.10 (https://doi.org/10.5281/zenodo.21235981) in paired-end mode and aligned to the modified reference genome using STAR v2.1.0 (92). The alignment index was built with --sjdbOverhang 149 to optimize alignment for 150 bp paired-end reads. The additional non-default STAR parameters were introduced: --quantMode GeneCounts -- twopassMode Basic --runRNGseed 0 --outFilterMultimapNmax 20 -- alignSJDBoverhangMin 1. Differential expression gene (DEG) analysis was performed in R v4.4.2 using DESeq2 (93). To identify expressed genes, only RNAs with a baseMean > 10 were retained. Only expressed RNAs with an FDR adjusted P-value < 0.05 were considered differentially expressed; no fold-change threshold was applied.

High-confidence C-to-U editing sites were identified using Bullseye (62). Aligned reads from each Dox-treated sample were compared against those from the corresponding untreated samples. Consistent with previous DART-seq analyses (62), editing sites were called at positions with a minimum of 10 unique reads, at least 2 mutations, an editing rate of 5-95, and at least a 1.25-fold increase in editing rate. The lower-bound threshold of 5% was selected to exclude false-positive editing signals observed in the absence of APOBEC1-YTH induction. This threshold falls within the range of thresholds used in previous studies (16,62,63). Only sites detected in at least two biological replicates and located within a RAC motif were retained. Sites located within introns or non-coding RNAs were excluded. The editing rate for each RNA was defined as the maximum editing rate among the called sites within the RNA, or within the RNA coding sequence (CDS) where specified.

Gene ontology (GO) enrichment analysis (94,95) was done using the clusterProfiler R package. For GO enrichment analysis, only expressed RNAs with an FDR adjusted P-value < 0.05, log2 fold change > 0.5 and a C-to-U editing rate > 5% were included.

### Analysis of PAR-CLIP data

PAR-CLIP data were obtained from Courtney et al. 2017 (50) (accession number: PRJNA383739). For use with clipplotr (v1.0.0), data were mapped to the GRCh38.p14 genome with GENCODE v49 basic genome annotation merged with Influenza A virus genome and annotation files. The goodwright/clipseq (v1.8) (https://github.com/goodwright/clipseq) pipeline, run with Nextflow (v25.04.2) (96), was used to trim adapters, quality control, map reads and call peaks. PCR deduplication was skipped. Prior to mapping to the genome using STAR, files were pre-mapped to a fasta file containing consensus sequences for small non-coding RNA, tRNA and ribosomal RNA. STAR parameters used: --outFilterMismatchNmax 20, -- outFilterMismatchNoverLmax 0.4. Crosslink position was taken as the middle of the read. Peak-calling was done using Clippy (https://github.com/ulelab/clippy) with default parameters. Visualisations used Clipplotr (v1.0.0), also using default parameters (--rollmean 10, --miniHeightAdjust 1.0, --minPromAdjust 1.0, -- minGeneCount 5) were used with clipplotr (v1.0.0) (60). Crosslink tracks were normalised by experiment library size, scaled to crosslinks per million and smoothed using the rolling mean of 50 nucleotides. For use with PEKA (v1.0.2) (59), data were solely mapped to the GRCh38.p14 genome with GENCODE v49 basic genome annotation and run using the Goodwright clipseq pipeline (v1.8) as described above. Cross-link bed files produced by Clippy from the pipeline using default parameters were used to analyse 6-mer enrichment using the PEKA (v1.0.2) package. Whole gene distribution and relative occurrence distribution files produced were used to plot relative occurrences heatmap.

## DATA AVAILABILITY

The DART-seq data reported in this paper is available at ArrayExpress (E-MTAB-17674). Any additional information required to reanalyse the data reported in this paper is available from the lead contact upon request.

## ACKNOWLEDGEMENTS

We thank members of the Neil, Holland and Swanson laboratories for helpful discussions. We also thank Charlotte Capitanchik for advice on analysing the PA-m6A-seq and PAR-CLIP datasets and Peter Hill for constructive comments on the manuscript. We thank Hannah Mischo, Rocio Martinez-Nunez and Wendy Barclay for reagents. We gratefully acknowledge the support of the HPC CREATE at King’s College London.

---

**Figure S1. Small molecule inhibition of METTL3 restricts IAV replication across different viral strain and airway epithelial cell lines.**

**(A)** A549 cells were pretreated for 2 h with STM2457 (60 μΜ) or an equivalent volume of vehicle control DMSO before infection with IAV A/California/7/2009 (H1N1) or IAV-PR8 at an MOI of 0.01. Infected cells were maintained in the presence of STM2457 or DMSO for 48 h. (B) HBEC3-KT cells were treated with STM2457 as described in Fig. 1D and infected with IAV-WSN at an MOI of 0.01 for 48 h. Viral titres were determined by plaque assay on MDCK cells. Data represent mean ± SD from n = 3 biological replicates. * denotes a statistically significant difference (p < 0.05), as determined by one-way ANOVA of log-transformed data. **(C)** Viability of HBEC3-KT cells following STM2457 treatment was assessed by flow cytometry using a live/dead cell stain.

**Figure S2. Motif enrichment analysis of PA-m^6^A-seq and YTHDF1-3 PAR-CLIP datasets from IAV-infected cells.**

Top 10 motifs ranked by PEKA score for PA-m6A-seq as well as YTHDF1, YTHDF2, YTHDF3 and GFP PAR-CLIP datasets.

**Figure S3. Distribution of m^6^A and YTHDF1-3 binding across IAV transcripts.**

PA-m^6^A-seq and YTHDF1, YTHDF2, YTHDF3 and GFP crosslinks were visualised across the IAV cRNAs.

**Figure S4. DART-seq enables quantitative analysis of m^6^A modifications in IAV-infected cells.**

**(A)** A549 cells stably expressing a YTH-APOBEC1 fusion protein with a C-terminal HA tag were generated by lentiviral transduction. Inducible expression of the fusion protein upon doxycycline treatment was confirmed by western blot. **(B)** Effect of YTH-APOBEC1 expression on IAV replication. Cells were infected with IAV-WSN at an MOI = 0.01 for 48 hours in the presence or absence of STM2457 (60 μΜ). APOBEC1-YTH expression was induced with 0.25 μg/mL doxycycline 24 hpi. Viral replication was assessed by plaque assay on MDBK cells. N = 3, mean ± SD. **(C)** Cumulative distribution of C-to-U editing frequencies in A549-YTH-APOBEC1 cells infected with IAV-WSN.

**Figure S5. DART-seq analysis of representative virus-induced genes.**

C-to-U editing frequencies across the RIPK1, ADAR, IFIT1, IFIT5, and TLR3 transcripts.

**Figure S6. PA-m^6^A-seq analysis of IFNB1 mRNA.**

PA-m^6^A-seq crosslinks were visualised across the IFNB1 transcript.

**Figure S7. PA-m^6^A-seq analysis of IFITM2 and IFITM3 transcripts.**

**(A-B)** PA-m^6^A-seq crosslinks were visualised across the (A) IFITM2 and (B) IFITM3 transcripts.

**Figure S8. NEDD4 knockdown or palmitoylation inhibition do not alter STM2457-mediated restriction of IAV replication.**

**(A)** Representative western blots from the cycloheximide chase experiments used to determine IFITM2/3 and MX1 protein half-lives shown in Fig. 6E-F. **(B)** Effect of siRNA-mediated NEDD4 depletion on STM2457-mediated restriction of IAV replication. **(C)** Effects of pharmacological inhibition of S-palmitoylation on IAV replication in the presence or absence of STM2457. Antibody B is 11714-I-AP.

**Figure S9. Cyclosporin H inhibits STM2457-mediated restriction of IAV replication.**

**(A)** Effect of cyclosporine H (CsH) treatment on IAV replication in the presence or absence of STM2457. **(B)** Effect of cyclosporine H (CsH) treatment on IFITM2/3 expression in the presence or absence of STM2457. Antibody B is 11714-I-AP.

