## Supplemental Figures and Tables for "N6-methyladenosine (m^6^A) RNA modifications control the antiviral response to influenza A virus infection by limiting IFITM2/3 expression"

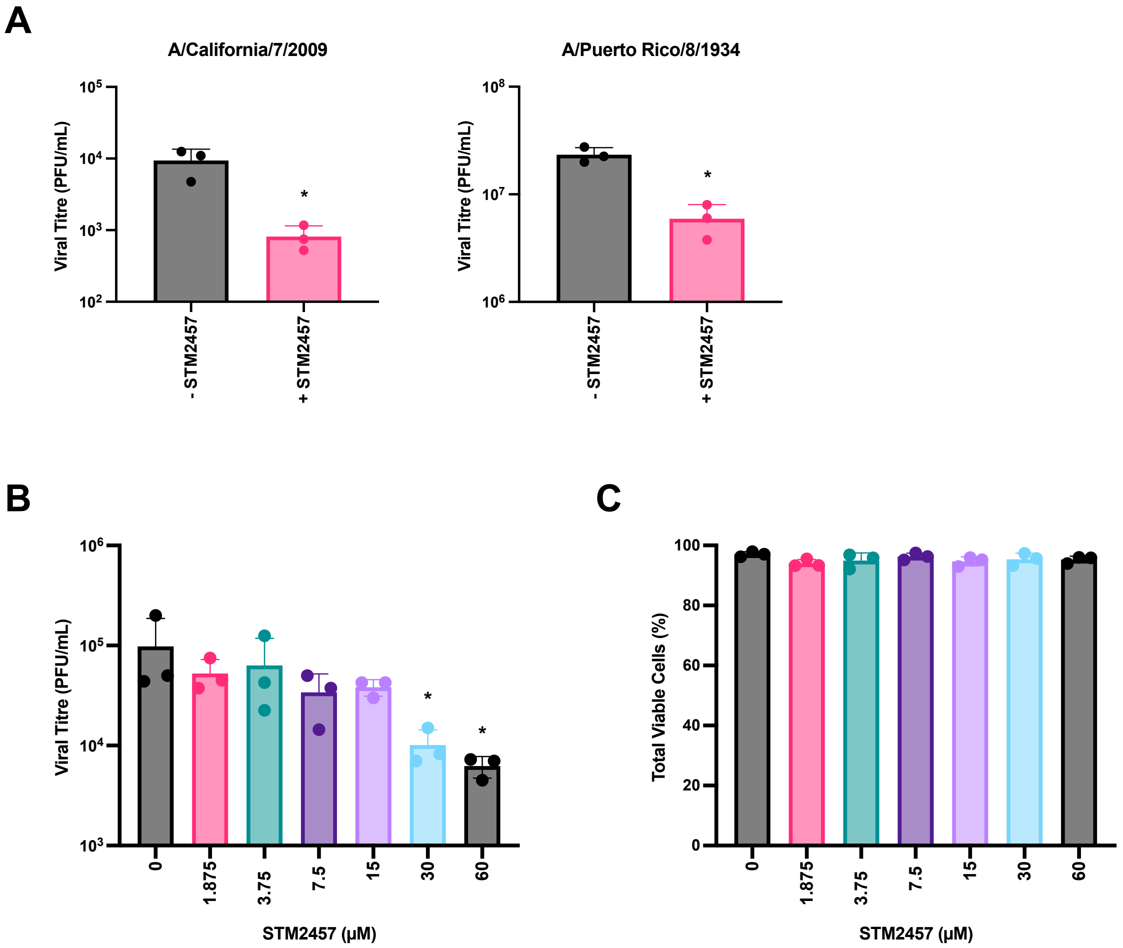
A
A/California/7/2009
Viral Titre (PFU/mL)
- STM2457
+ STM2457
A/Puerto Rico/8/1934
B
Viral Titre (PFU/mL)
STM2457 (µM)
C
Total Viable Cells (%)
STM2457 (µM)

Figure S1

**A**

| m6A motifs | PEKA score |
|---|---|
| GAGGAC | 57.0 |
| GGACUG | 53.9 |
| GACUGG | 46.7 |
| CAGGAC | 46.6 |
| GGGCUG | 41.5 |
| CUGGAC | 41.5 |
| AGGACU | 40.8 |
| GGACAG | 38.9 |
| CGGACC | 36.9 |
| GGACUC | 36.5 |

**B**

| YTHDF1 motif | PEKA-score |
|---|---|
| CUGGAC | 32.2 |
| CCGGAC | 29.0 |
| CAGACU | 26.5 |
| CGGACC | 23.7 |
| UGGACC | 22.2 |
| GGACCU | 21.1 |
| UGGACA | 20.1 |
| GACUGU | 19.7 |
| CGGACA | 18.7 |
| CGGACU | 17.9 |

**C**

| YTHDF2 motif | PEKA-score |
|---|---|
| CUGGAC | 15.7 |
| CCGGAC | 14.7 |
| AUGCGU | 12.2 |
| CGGACC | 12.1 |
| GGACUU | 12.0 |
| CGUGAC | 12.0 |
| AGGACU | 12.0 |
| GGACUC | 11.7 |
| GACGAU | 11.1 |
| UACGGA | 11.1 |

**D**

| YTHDF3 motif | PEKA-score |
|---|---|
| UACGCU | 11.2 |
| GGACGA | 11.1 |
| UGGCCG | 10.9 |
| UGCGCA | 10.6 |
| CAGGAC | 10.5 |
| CUGGAC | 10.1 |
| AUCCCG | 10.0 |
| UGUAAA | 9.9 |
| AAACGU | 9.8 |
| UCUACG | 9.7 |

**E**

| GFP motifs | PEKA score |
|---|---|
| AAACGC | 12.0 |
| CAAACG | 10.4 |
| UAACGU | 10.3 |
| UACACG | 10.1 |
| AUCGAU | 10.0 |
| ACUAUC | 9.9 |
| UAUGCG | 9.7 |
| CUACGG | 9.6 |
| CGGUAA | 9.5 |
| ACGCGU | 9.5 |

**Figure S2**

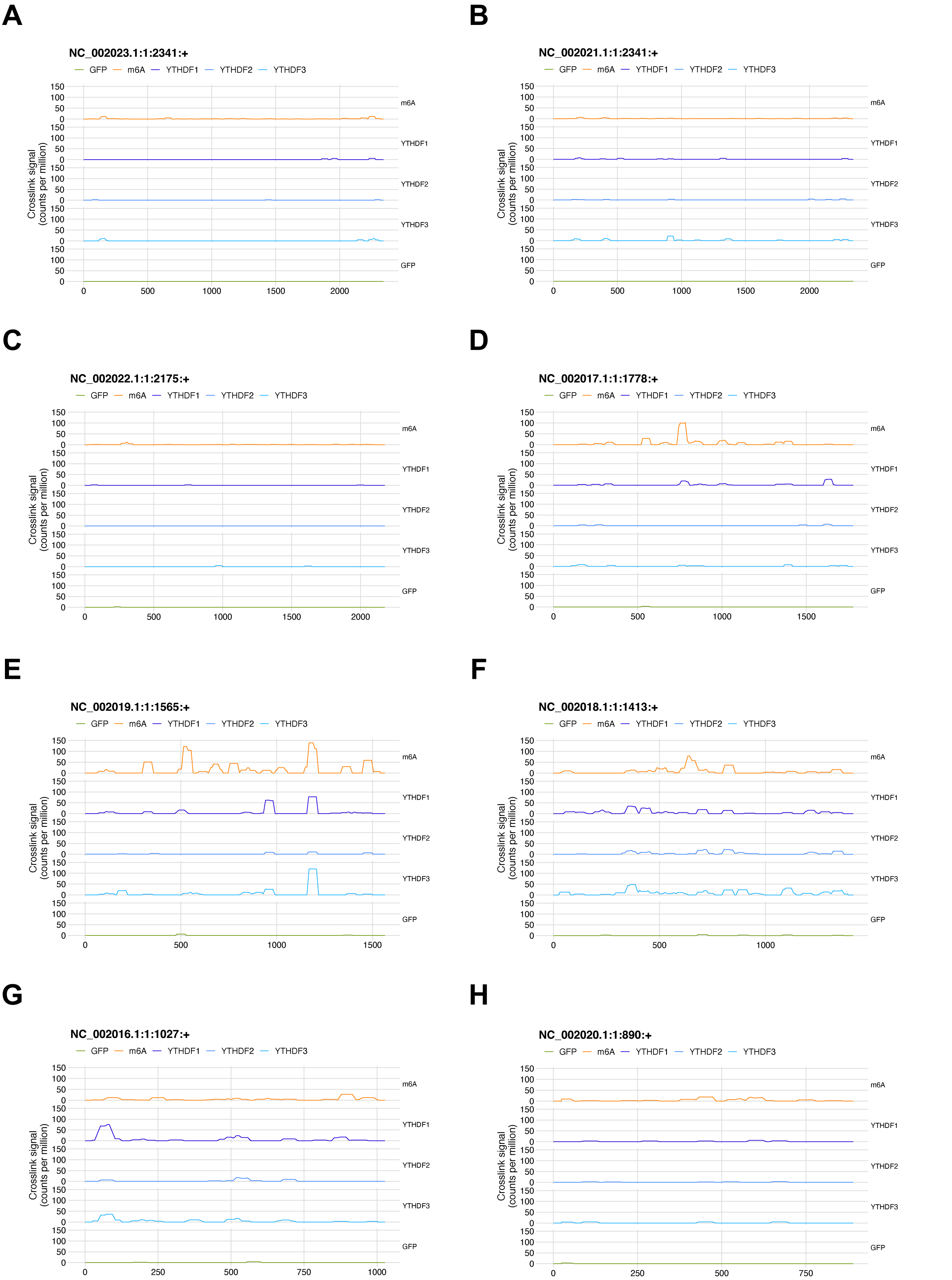
A
NC_002023.1:1:2341:+
B
NC_002021.1:1:2341:+
C
NC_002022.1:1:2175:+
D
NC_002017.1:1:1778:+
E
NC_002019.1:1:1565:+
F
NC_002018.1:1:1413:+
G
NC_002016.1:1:1027:+
H
NC_002020.1:1:890:+
GFP
m6A
YTHDF1
YTHDF2
YTHDF3
Crosslink signal (counts per million)
0
50
100
150
500
1000
1500
2000
250
750

Figure S3

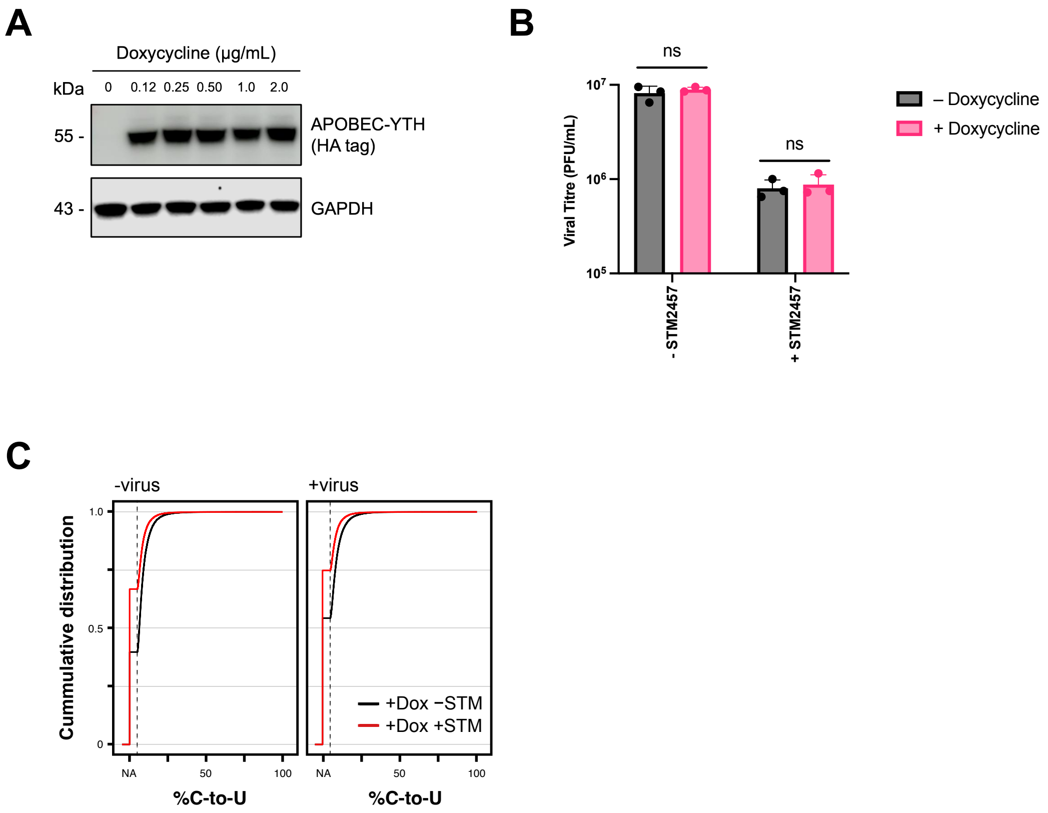
A
Doxycycline (µg/mL)
kDa
0
0.12
0.25
0.50
1.0
2.0
55 -
APOBEC-YTH
(HA tag)
43 -
GAPDH
B
ns
ns
Viral Titre (PFU/mL)
10^7
10^6
10^5
- STM2457
+ STM2457
– Doxycycline
+ Doxycycline
C
-virus
+virus
Cummulative distribution
1.0
0.5
0
NA
50
100
%C-to-U
+Dox −STM
+Dox +STM

Figure S4

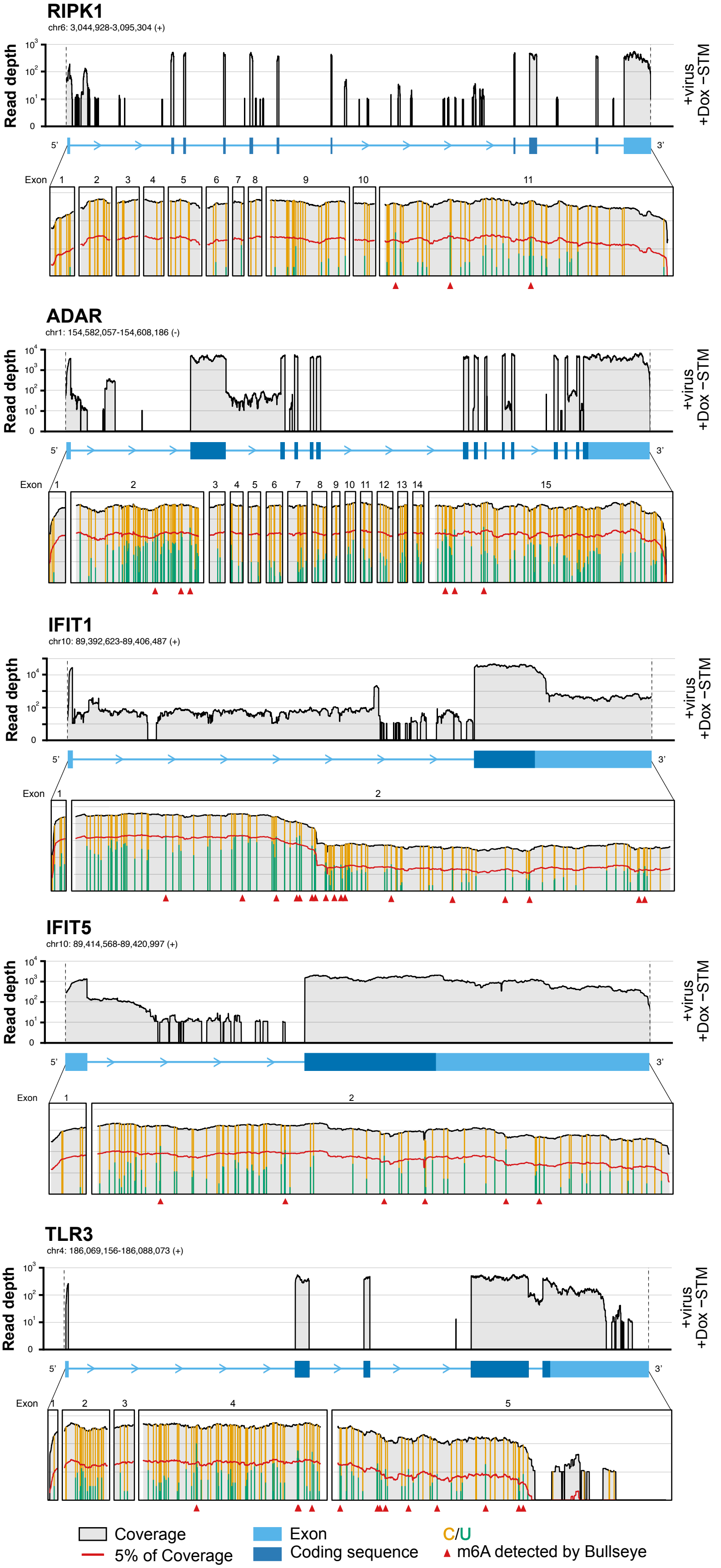

RIPK1
chr6: 3,044,928-3,095,304 (+)
ADAR
chr1: 154,582,057-154,608,186 (-)
IFIT1
chr10: 89,392,623-89,406,487 (+)
IFIT5
chr10: 89,414,568-89,420,997 (+)
TLR3
chr4: 186,069,156-186,088,073 (+)
Read depth
+virus
+Dox −STM
Exon
5'
3'
Coverage
5% of Coverage
Exon
Coding sequence
C/U
m6A detected by Bullseye


Figure S5

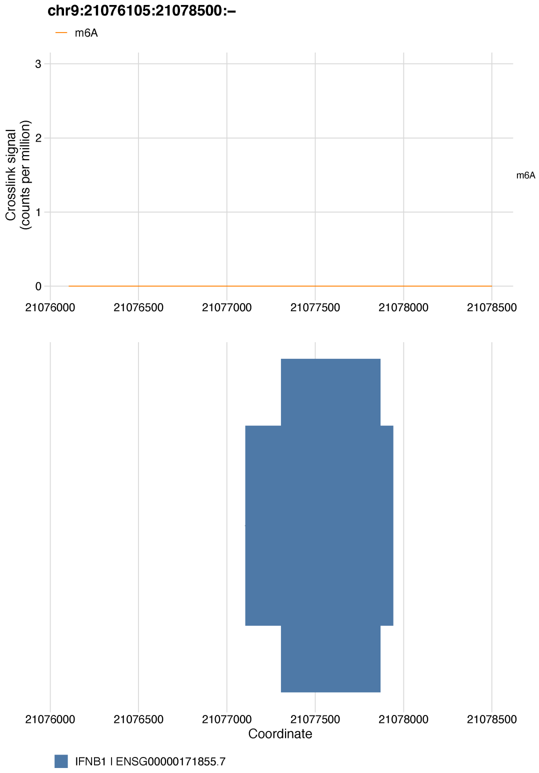
chr9:21076105:21078500:–
m6A
Crosslink signal
(counts per million)
3
2
1
0
21076000
21076500
21077000
21077500
21078000
21078500
m6A
Coordinate
IFNB1 I ENSG00000171855.7

**Figure S6**

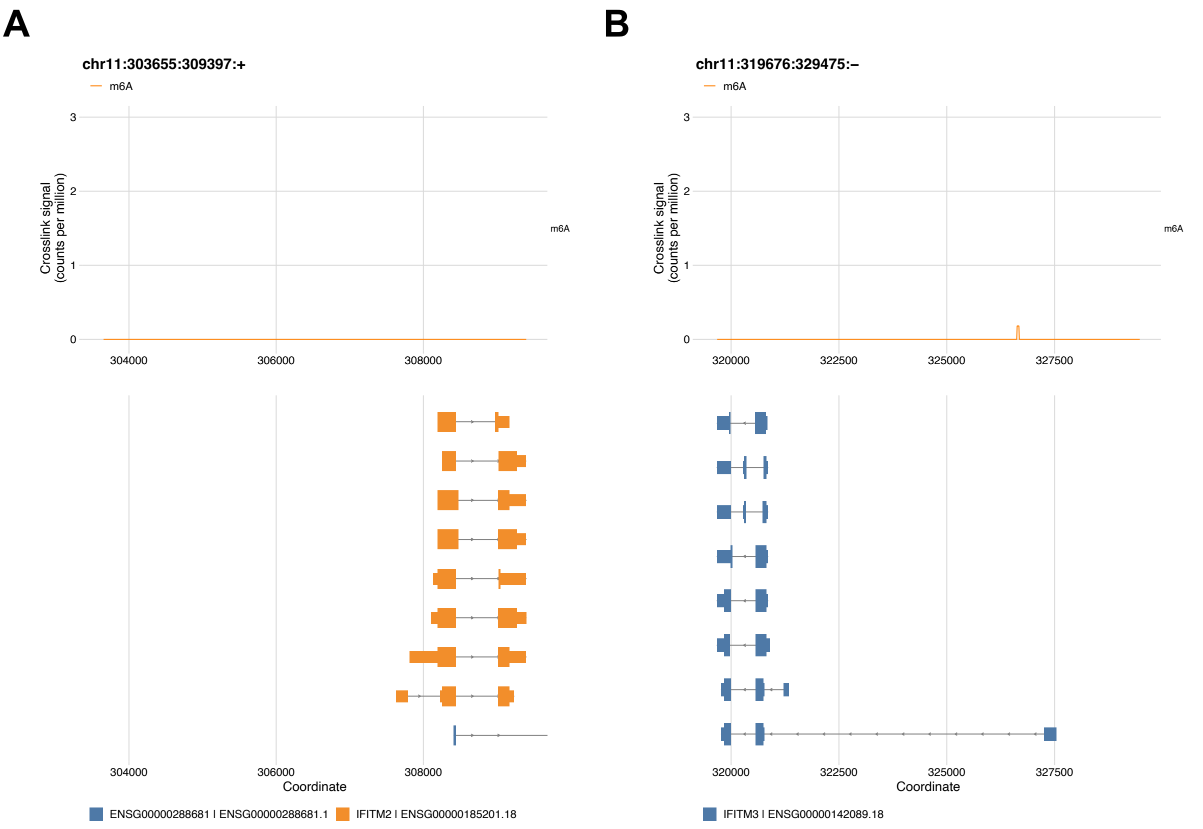

A
chr11:303655:309397:+
m6A
Crosslink signal (counts per million)
0
1
2
3
304000
306000
308000
Coordinate
ENSG00000288681 | ENSG00000288681.1
IFITM2 | ENSG00000185201.18
B
chr11:319676:329475:–
m6A
Crosslink signal (counts per million)
320000
322500
325000
327500
Coordinate
IFITM3 | ENSG00000142089.18


**Figure S7**

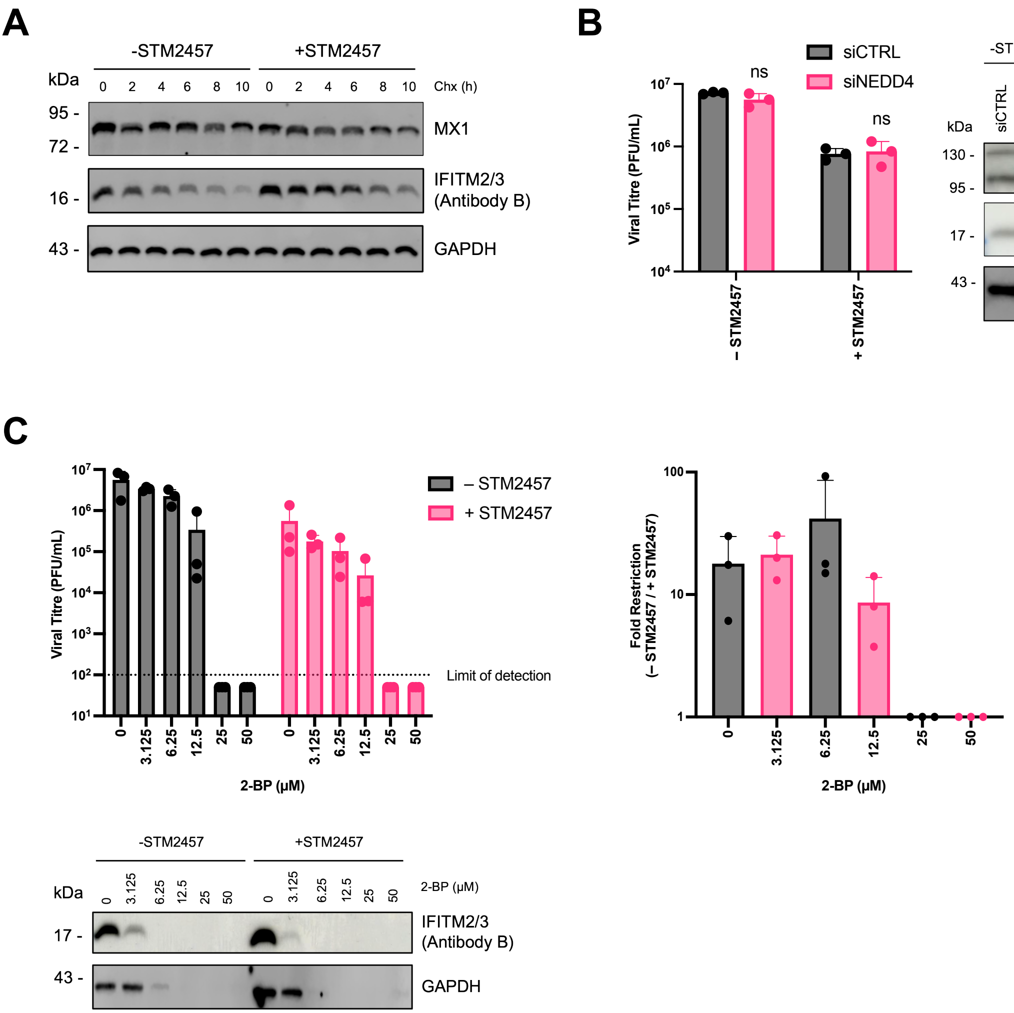
A
-STM2457
+STM2457
kDa
0 2 4 6 8 10 0 2 4 6 8 10 Chx (h)
95 -
72 -
MX1
16 -
IFITM2/3
(Antibody B)
43 -
GAPDH
B
siCTRL
siNEDD4
ns
ns
Viral Titre (PFU/mL)
– STM2457
+ STM2457
-STM2457
+STM2457
kDa
siCTRL
siNEDD4
siCTRL
siNEDD4
130 -
95 -
NEDD4
17 -
IFITM2/3
(Antibody B)
43 -
GAPDH
C
– STM2457
+ STM2457
Viral Titre (PFU/mL)
Limit of detection
0 3.125 6.25 12.5 25 50 0 3.125 6.25 12.5 25 50
2-BP (μM)
Fold Restriction
(– STM2457 / + STM2457)
100
10
1
0 3.125 6.25 12.5 25 50
2-BP (μM)
-STM2457
+STM2457
kDa
0 3.125 6.25 12.5 25 50 0 3.125 6.25 12.5 25 50 2-BP (μM)
17 -
IFITM2/3
(Antibody B)
43 -
GAPDH

Figure S8

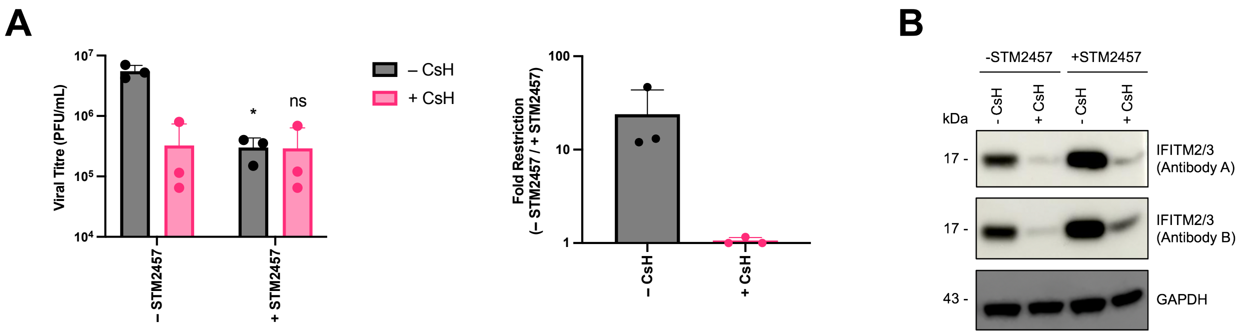
A
Viral Titre (PFU/mL)
10^7
10^6
10^5
10^4
*
ns
– STM2457
+ STM2457
– CsH
+ CsH
Fold Restriction
(– STM2457 / + STM2457)
100
10
1
– CsH
+ CsH
B
-STM2457
+STM2457
- CsH
+ CsH
- CsH
+ CsH
kDa
17 -
17 -
43 -
IFITM2/3
(Antibody A)
IFITM2/3
(Antibody B)
GAPDH

**Figure S9**

**Table S2: siRNA reagents**

| TARGET | SOURCE | CAT. # |
|---|---|---|
| Non-targeting CTRL | Dharmacon | D-001210-04-05 |
| METTL3 | Dharmacon | M-005170-01-0005 |
| METTL14 | Dharmacon | M-014169-00-0005 |
| YTHDC1 | Dharmacon | M-015332-01-0005 |
| YTHDF1 | Dharmacon | M-018095-01-0005 |
| YTHDF2 | Dharmacon | M-021009-01-0005 |
| YTHDF3 | Dharmacon | M-017080-01-000 |
| IFITM2 | Dharmacon | M-020103-02-0010 |
| IFITM3 | Dharmacon | L-014116-01-0010 |
| NEDD4 | Dharmacon | M-007178-02-0005 |

**Table S3: Antibodies**

| ANTIBODY | SPECIES | DILUTION | SOURCE | CAT. # |
|---|---|---|---|---|
| **Primary Antibody** | | | | |
| Anti-METTL3 | Rabbit | 1:1000 | Proteintech | 15073-1-AP |
| Anti-METTL14 | Rabbit | 1:1000 | Proteintech | 26158-1-AP |
| Anti-YTHDC1 | Rabbit | 1:1000 | Atlas Antibodies | HPA036462 |
| Anti-YTHDF1 | Rabbit | 1:1000 | Proteintech | 17479-1-AP |
| Anti-YTHDF2 | Rabbit | 1:2000 | Proteintech | 24744-1-AP |
| Anti-YTHDF3 | Rabbit | 1:5000 | Proteintech | 25537-1-AP |
| Anti-ALKBH5 | Rabbit | 1:5000 | Proteintech | 16837-1-AP |
| Anti-HA tag | Rabbit | 1:1000 | Cell Signaling Tech. | 3724 |
| Anti-GAPDH | Mouse | 1:2000 | Proteintech | 60004-1-Ig |
| Anti-RIG-I | Rabbit | 1:1000 | Enzo Life Sciences | AT111 |
| Anti-MDA5 | Rabbit | 1:1000 | Cell Signaling Tech. | D74E4 |
| Anti-MAVS | Rabbit | 1:2000 | Bethyl | A300-782A |
| Anti-MX1 | Rabbit | 1:1000 | Genetex | GTX111153 |
| Anti-NCOA7(AS) | Rabbit | 1:500 | Prof. Michael Malim | N/A |
| Anti-TRIM25 | Rabbit | 1:2500 | Abcam | ab167154 |
| Anti-IFITM1 | Mouse | 1:3000 | Proteintech | 60074-1-Ig |
| Anti-IFITM2/3 (Antibody A) | Rabbit | 1:3000 | Proteintech | 12769-1-AP |
| Anti-IFITM2/3 (Antibody B) | Rabbit | 1:3000 | Proteintech | 11714-1-AP |
| Anti-NP | Rabbit | 1:5000 | Genetex | GTX125989 |
| Anti-NEDD4 | Rabbit | 1:1000 | Proteintech | 21698-1-AP |

| **Secondary Antibody** | | | | |
|---|---|---|---|---|
| Anti-rabbit | Goat | 1:10000 | Li-COR | 926-32211 |
| Anti-mouse | Goat | 1:10000 | Li-COR | 925-68070 |
| Anti-rabbit HRP | Goat | 1:5000 | Cell Signaling Tech. | 7074 |
| Anti-mouse HRP | Horse | 1:5000 | Cell Signalling Tech. | 7076 |

**Table S4: RT-PCR assays**

| GENE | ASSAY ID |
|---|---|
| IFNB1 | Hs01077958_s1 |
| IFITM2 | Hs00829485_sH |
| IFITM3 | Hs03057129_s1 |
| 18s rRNA | Hs99999901_s1 |
